# Analysis of Infected Host Gene Expression Reveals Repurposed Drug Candidates and Time-Dependent Host Response Dynamics for COVID-19

**DOI:** 10.1101/2020.04.07.030734

**Authors:** Jing Xing, Rama Shankar, Aleksandra Drelich, Shreya Paithankar, Evgenii Chekalin, Thomas Dexheimer, Mei-Sze Chua, Surender Rajasekaran, Chien-Te Kent Tseng, Bin Chen

**Author notes:** Correspondence to Bin Chen, Kent Chien-Te, Tseng. These authors contributed equally (Jing Xing, Rama Shankar, Aleksandra Drelich).

## Abstract

The repurposing of existing drugs offers the potential to expedite therapeutic discovery against the current COVID-19 pandemic caused by the SARS-CoV-2 virus. We have developed an integrative approach to predict repurposed drug candidates that can reverse SARS-CoV-2-induced gene expression in host cells, and evaluate their efficacy against SARS-CoV-2 infection *in vitro*. We found that 13 virus-induced gene expression signatures computed from various viral preclinical models could be reversed by compounds previously identified to be effective against SARS- or MERS-CoV, as well as drug candidates recently reported to be efficacious against SARS-CoV-2. Based on the ability of candidate drugs to reverse these 13 infection signatures, as well as other clinical criteria, we identified 10 novel candidates. The four drugs bortezomib, dactolisib, alvocidib, and methotrexate inhibited SARS-CoV-2 infection-induced cytopathic effect in Vero E6 cells at < 1 µM, but only methotrexate did not exhibit unfavorable cytotoxicity. Although further improvement of cytotoxicity prediction and bench testing is required, our computational approach has the potential to rapidly and rationally identify repurposed drug candidates against SARS-CoV-2. The analysis of signature genes induced by SARS-CoV-2 also revealed interesting time-dependent host response dynamics and critical pathways for therapeutic interventions (e.g. Rho GTPase activation and cytokine signaling suppression).

## Introduction

Since early December 2019, the newly emerged SARS-CoV-2 has infected more than 4 million people globally (COVID-19 Report from WHO n.d.). In the United States, confirmed cases increased exponentially and steadily, exceeding the million mark within two months, making the United States the hardest hit country globally (CDC Cases and Updates n.d.). Among all infected patients, approximately 16% suffered from severe acute respiratory distress syndrome (ARDS) and approximately 6% died from acute respiratory failure, acute cardiac injury, secondary infection, and other serious complications (COVID-19 Report from WHO n.d.; Huang *et al*. 2020). As this highly contagious and pathogenic virus spread rapidly throughout the world, the WHO declared it a pandemic (named COVID-19), and effective therapeutics are urgently needed. Repurposing existing drugs could be an efficient and timely means of identifying novel drugs for treating this disease. Initially, repurposed drugs such as lopinavir/ritonavir, baricitinib, remdesivir, and chloroquine were investigated for clinical efficacy in COVID-19 patients (Peeri *et al*. 2020; Wang *et al*. 2020). Remdesivir has since shown encouraging efficacy, and is currently used as a benchmark drug for treating COVID-19 patients with moderate to severe symptoms. These repurposed drugs under study are hypothesized to target key steps of viral entry, or specific proteins involved in viral replication, including viral proteases (Zumla *et al*. 2016). In addition to viral replication, the viral pathogen associated molecular pattern (PAMP) (e.g., immune dysfunction and endoplasmic reticulum stress, Figure 1A) could be targeted to improve clinical outcomes (de Wit *et al*. 2016). PAMPs-mediated signaling pathways are attractive drug targets in diseases caused by human pathogens. Therefore, effectively targeting these pathways to stop the progression to acute respiratory distress syndrome caused by SARS-CoV-2 might potentially reduce the morbidities and mortalities associated with this virus. Independent of SARS-CoV-2 infection, in aging adult populations, ARDS is associated with mortality rates of 30-50% (Bellani *et al*. 2016). Thus, a methodical and unbiased search for new drug candidates from a large drug library could uncover drugs that have the potential to arrest the infection and ameliorate its debilitating effects. We aim to accomplish this by analyzing infection-induced gene expressions in the host cells.

**Figure 1.**
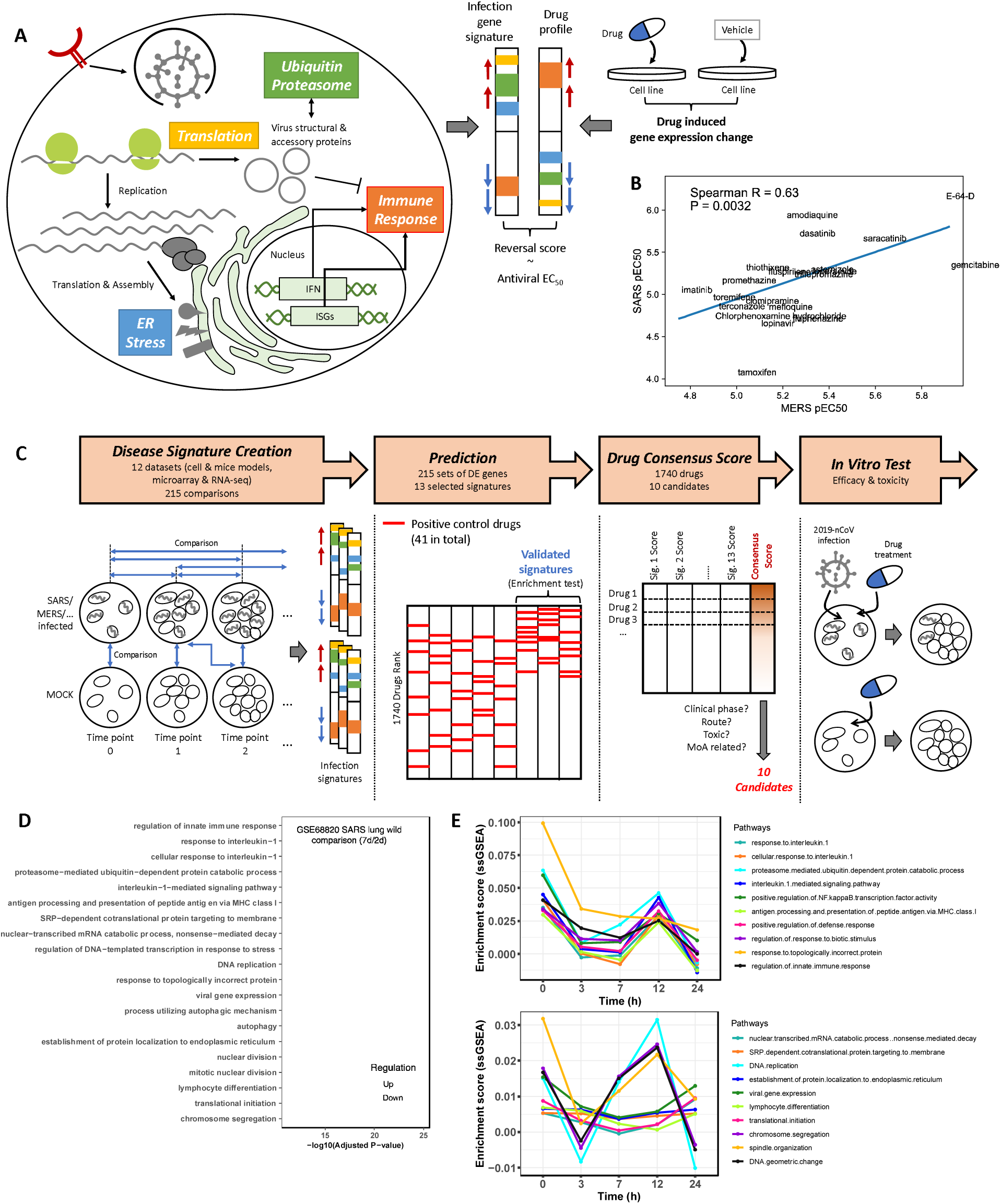
Study workflow and biological processes dysregulated by viral infection. **A**, An illustration of reversing the expression of host genes comprising multiple biological processes (highlighted with colors) induced by coronavirus infection. Drug-induced gene expression profiles are taken from the LINCS database. A good candidate should activate the repressed biological processes and inhibit the upregulated processes. **B**, Correlation of the published antiviral activities of 30 drugs (pEC_50_, -log_10_ transformed EC_50_ value in mol/L) against MERS-CoV and SARS-CoV. **C**, Study workflow including creation of disease signatures, prediction of drug candidates, selection of a final drug list, and *in vitro* validation. One disease signature composed of the differentially expressed genes of each comparison led to one drug prediction list. Only the signature resulting in a prediction list where known positive drugs were enriched on the top was considered a valid signature. **D**, Dysregulated pathways in lungs after SARS-CoV infection at 7 day compared with 2 day. **E**, Dynamics of dysregulated pathways (Top: Downregulated pathways, and Down: upregulated pathways) in MDC001 cells infected with MERS-CoV across different post-infection times (0 h-24 h) (study id: GSE79172). Only one study was selected for **D** and **E** each; the dysregulated pathways and their dynamics for other studies are available in supplementary materials (Figure S2 and S3, Extended Data 1).

We and others (B. Chen, Wei, *et al*. 2017; Lee *et al*. 2017; van Noort *et al*. 2014; Pessetto *et al*. 2016; Qu and Rajpal 2012; Sirota *et al*. 2011; H. Wu *et al*. 2017) have utilized a systems-based approach that employs gene expression profiles of disease samples and drug-induced gene expression profiles from cell lines to discover new therapeutic candidates for diseases. The central idea is to identify drugs that reverse the disease gene expression signature by suppressing the over-expressed disease genes and activating the repressed genes (Figure 1A). A disease signature is defined as a list of differentially expressed genes between disease samples and normal control samples. We recently established a scoring system for the reversal of gene expression (named summarized Reversal of Gene Expression Score, sRGES) that correlates with drug efficacy in cancers (B. Chen, Ma, *et al*. 2017), demonstrating the feasibility of applying this approach to predict drug candidates for other diseases, including COVID-19 and other viral infections.

## Results

### Creating and Validating Host Infection Signature

To utilize sRGES for drug discovery in SARS-CoV-2, we first needed to collect SARS-CoV-2-induced host gene expression profiles, which were not available at the time of writing. Given the high genomic similarity of SARS-CoV-2 with SARS-CoV (79%), and with MERS-CoV (50%) (H, Wang, *et al*. 2020) (Lu et al. 2020), we reasoned that existing host gene expression profiles of samples infected by SARS-CoV or MERS-CoV could approximate to those infected by SARS-CoV-2. To verify this assumption, we compiled 331 signatures of various viruses from enrichR and GEO (Table S1) and used an established pipeline to score 1740 drugs in our repurposing library for their ability to reverse virus-induced signatures (B. Chen, Ma, et al. 2017; Zeng et al. 2019). Clustering of these signatures based on their drug prediction scores suggests that signatures derived from the same virus or virus family under similar experimental conditions tend to cluster together (Figure S1). An example cluster includes one signature derived from primary human microvascular endothelial cells (MMVE001) after two days of MERS-CoV infection (study id: GSE79218), and another derived from melanoma cells in mice after seven days of SARS-CoV infection (study id: GSE68820). In addition, Spearman correlation coefficient of the *in vitro* drug efficacy data (EC_50_: Half maximal effective concentration) for the SARS-CoV and MERS-CoV datasets is close to 0.6 (Figure 1B). The clustering and correlation results suggested that drugs predicted based on the signatures induced by SARS-CoV or MERS-CoV may have potential applicability in SARS-CoV-2. Therefore, we developed a pipeline to identify repurposed drugs against MERS-CoV and SARS-CoV, and then experimentally evaluate these drugs in SARS-CoV-2 (Figure 1C).

In total, 430 samples from public repositories, representing infection by MERS-CoV or SARS-CoV (and a few other strains for comparison) in different models (e.g., cell line, mouse models) across multiple time points were used to identify disease signatures (Table S1, 12 studies in total). Their expression profiles were created using either microarray or RNA-Sequencing. Depending on the profiling platform, data processing and signature creation methods varied (see Methods). The previous clusters (Figure S1) were highly confounded by post-infection time points (Figure S1), meaning the disease signatures and their predicted drugs were strikingly different under different time points. Therefore, we enumerated all the possible comparisons (Figure 1C), including (1) comparisons between infection and mock groups at each time point, (2) comparisons between different time points within each of the infection or the mock group (e.g., time point 1 *vs*. time point 0, time point 2 *vs*. time point 1), and (3) comparisons both between time points and between infection and mock groups. These comparisons revealed different virus-related biological processes and their dynamic regulation. For instance, analysis of SARS-CoV infected lung tissue data showed that various biological processes, including viral gene expression, DNA replication, nuclear division, lymphocytes differentiation and translation-related processes, were activated (Figure 1D, S2 and Extended Data 1). In contrast, interleukin and autophagy-related processes were repressed in infected samples (Figure 1D, S2 and Extended Data 1). Interestingly, these processes displayed time-dependent dynamics in infected samples (e.g., 3 h and 12 h in Figure 1E). More examples of infection dynamics from other studies are shown in Figure S3. For instance, immune signaling pathways were down-regulated dramatically within the first 3 hours after infection, while DNA replication related pathways were first suppressed during this period but then activated until 12 h post-infection. These observed host dynamic responses to virus infection suggests that comparisons between different time points are important to capture representative biological events throughout the course of a viral infection.

For each comparison, we computed a disease signature to characterize the infection status, followed by the prediction of which drugs may have activity (ability to reverse disease signature). As we could not directly evaluate the quality and pathologic relevance of each disease signature, we validated them using those drugs identified to be positive in *in vitro* MERS/SARS-CoV testing (41 positive drugs in total, 30 with EC_50_ values, Table S2). Among 215 MERS-CoV or SARS-CoV infection signatures, only 13 signatures were able to recover these positive drugs (which were shown to be highly enriched at the top of the predicted drug lists) (Extended Data 2 and 3). Moreover, EC_50_ of these drugs significantly correlated with sRGES (Figure 2A and Figure S4). Validating our analysis, we did not observe significant enrichment of anti-coronavirus positive drugs using H1N1 infection signatures (Extended Data 2). Recent clinical data suggested that the combination of lopinavir/ritonavir offers clinical benefits to patients infected with SARS-CoV-2 (Chu *et al*. 2004; Lim *et al*. 2020). Both drugs were profiled in LINCS (Subramanian et al. 2017), the drug library we primarily used. individually; however, none of them alone could induce substantial gene expression changes (absolute z score < 2) under 10 µM concentration in cancer cell lines; therefore, neither was predicted as a hit. We next found another dataset where HepaRG cells were treated with ritonavir at 10 different concentrations (ranging from 9 nM to 300 µM). We observed that ritonavir could reverse nearly all the 13 disease signatures under high concentrations (between 10 µM to 200 µM) (Figure S5). Thus, we reasoned that the top ranked drugs with highly negative sRGES in these valid comparisons could be new therapeutic candidates.

**Figure 2.**
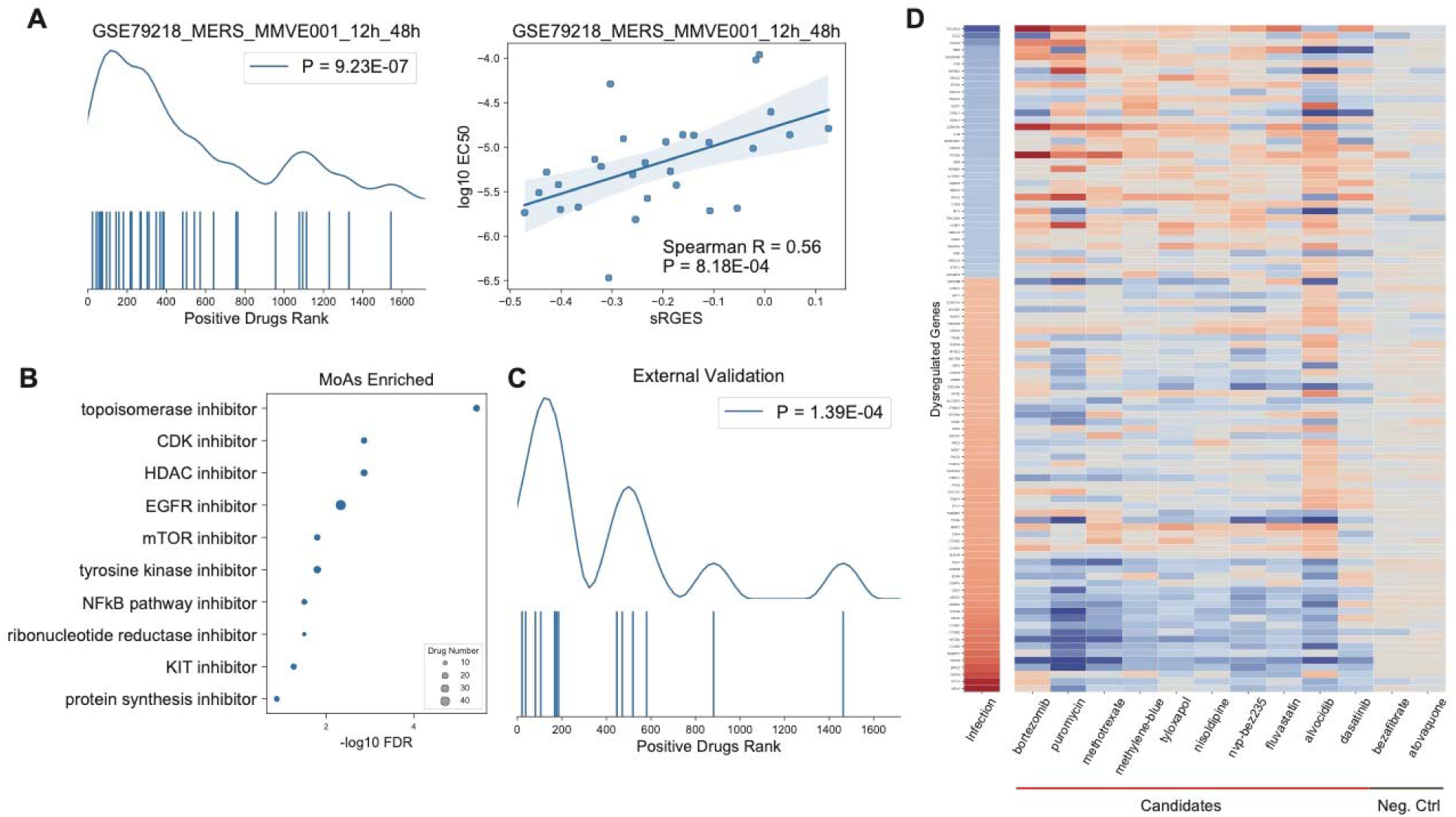
Meta drug analysis and candidate evaluation. **A**, An example of a validated signature derived from differentially expressed genes between 12 h and 48 h after MERS-CoV infection in the MMVE001 cell line (GSE79218 dataset). The left panel shows the enrichment of positive drugs (p value was computed by Wilcoxon rank sums test, see Methods). The curve shows enrichment density, and each bar under the curve represents the rank of a positive drug on the prediction list. The right panel shows the correlation between sRGES and EC_50_ (in mol/L, log_10_ transformed) of the positive controls. Each point indicates a positive drug. **B**, Enriched MoA (blue dots) and their FDR based on consensus drug ranking. Dot size corresponds to the number of drugs associated with the MoA. **C**, External validation using published anti-SARS-CoV-2 efficacy of FDA-approved drugs (Jeon *et al*. 2020). **D**, Heatmap of the summarized infection signature and the summarized reversal effects of 10 selected candidates and two random examples (see Methods). Red color indicates up-regulated genes and blue indicates down-regulated genes.

### Consensus Scoring for Final Drug Prediction

Merging disease signatures derived from different platforms and biological conditions is highly complicated; therefore, instead of merging them, we calculated the consensus score of all the drug prediction lists derived from individual valid disease signatures. We took the median of the ranks across multiple comparisons for each drug, and ranked them based on their median rank (Extended Data 4). Mechanism of action (MoA) enrichment analysis of the final ranked drug list revealed a number of significant drug classes including CDK inhibitors, mTOR inhibitors, and NF-κB pathway inhibitors (Figure 2B). Recently, Jeon *et al*. (Jeon *et al*. 2020) tested 35 FDA-approved drugs for potential activity against SARS-CoV-2, among which 14 positive drugs were also represented in our screening library. These drugs were also significantly enriched at the top of our final prediction (Figure 2C, p = 1.39E-4), further validating our analysis. Interestingly, three anti-parasitic drugs pyrvinium, ivermectin, and niclosamide were ranked among the top 30 predicted drugs. Both ivermectin and niclosamide were shown to inhibit the replication of SARS-CoV-2 *in vitro* (Caly *et al*. 2020; Jeon *et al*. 2020), and pyrvinium and niclosamide were shown to be effective against MERS-CoV and SARS-CoV (Shen *et al*. 2019; C.-J. Wu *et al*. 2004).

### Selection and *In Vitro* Validation of Drug Candidates

We manually inspected the top candidates and selected ten representative candidates according to their reversal scores (Figure 2D, Extended Data 4), clinical applicability, MoA, and administration routes (Table S3). We then evaluated their cytotoxicity and ability to prevent cytopathic effect (CPE) in the Vero E6 cell line (Table 1). Instead of measuring EC_50_, we used more stringent criteria, which required a 4-day complete prevention of CPE observed under the microscope. Seven of the proposed drugs could completely prevent CPE at concentrations < 10 µM. However, five of these presented unfavorable cytotoxicity at their effective concentration, except methotrexate. In a repeat experiment (using a shorter drug treatment time of 2 h instead of 3 h), we examined the efficacy and toxicity for the four most effective drugs (preventing CPE at concentrations ≤ 1 µM): bortezomib, methotrexate, nvp-bez235, and alvocidib, as well as chloroquine (one of the drugs being tested clinically for SARS-CoV-2). However, under this shorter drug treatment time, none of the predicted drugs could completely prevent CPE as effectively as chloroquine (which did so at 15 µM).

**Table 1.** Efficacy and toxicity of selected predicted drugs against SARS-CoV-2. Since none of these predicted drugs are more effective than chloroquine, no further validations were conducted beyond this. Chloroquine was not included in the first round as it was not known as an effective hit when the experiment started.

| | MoA | Primary Indication | CPE Preventing<sup>*1</sup> ($\mu\text{M}$) | Toxic<sup>*2</sup> ($\mu\text{M}$) | Repeat<sup>*3</sup> CPE Preventing ($\mu\text{M}$) | Repeat Toxic ($\mu\text{M}$) |
| --- | --- | --- | --- | --- | --- | --- |
| Bortezomib | Proteasome inhibitor | Multiple myeloma | 0.05 | 0.002 | > 30 | > 30 |
| Puromycin | Protein synthesis inhibitor | Antibiotic | 20 | 0.002 | -<sup>*4</sup> | - |
| Methotrexate | Dihydrofolate reductase inhibitor | Rheumatoid arthritis | 0.78 | > 25 | > 30 | 30 |
| Methylene-blue | Guanylyl cyclase inhibitor | Methemoglobinemia | 20 | 5 | - | - |
| Tyloxapol | NF-kB pathway inhibitor | Bronchopulmonary secretions with mucus and pus | > 100 | 20 | - | - |
| Nisoldipine | Calcium channel blocker | Hypertension | 6.25 | 0.2 | - | - |
| Nvp-bez235 | PI3K/MTOR inhibitor | Advanced Solid Malignancies | 1 | 1 | > 30 | 7.5 |
| Fluvastatin | HMGCR inhibitor | Hypercholesterolemia | 3.125 | 0.2 | - | - |
| Alvocidib | CDK inhibitor | Acute myeloid leukemia | 0.05 | 0.002 | > 30 | 3.75 |

| | MoA | Primary Indication | CPE Preventing<sup>*1</sup><br>(μM) | Toxic<sup>*2</sup><br>(μM) | Repeat<sup>*3</sup> CPE Preventing<br>(μM) | Repeat Toxic (μM) |
| --- | --- | --- | --- | --- | --- | --- |
| Dasatinib | SRC/ABL kinase inhibitor | Leukemia | 3.125 | 1.6 | - | - |
| Chloroquine<sup>*5</sup> | Endosomal acidification inhibitor | Malaria | - | - | 15 | > 30 |

<sup>\*1</sup>, the lowest concentration of a drug to prevent CPE.
<sup>\*2</sup>, the lowest concentration of a drug showing cytotoxicity.
<sup>\*3</sup>, when repeated, drug treatment time was shorter (2 hours) than the previous test (3 hours).
<sup>\*4</sup>, not tested.
<sup>\*5</sup>, positive control of this assay.

### Enrichment analysis and time-dependent host response dynamics suggest therapeutic intervention points

The *in silico* external validations and our *in vitro* validations supported the robustness of our valid 13 infection signatures for drug discovery against pathogenic coronaviruses including SARS-CoV-2. To understand the underlying biology of these signatures and the impact of coronaviruses on the host biology, we performed canonical pathway and gene ontology enrichment analysis of the commonly dysregulated genes (i.e., up- or down-regulated in more than half of the 13 infection signatures, Figure 3, Extended Data 3 and Table S4).

**Figure 3.**
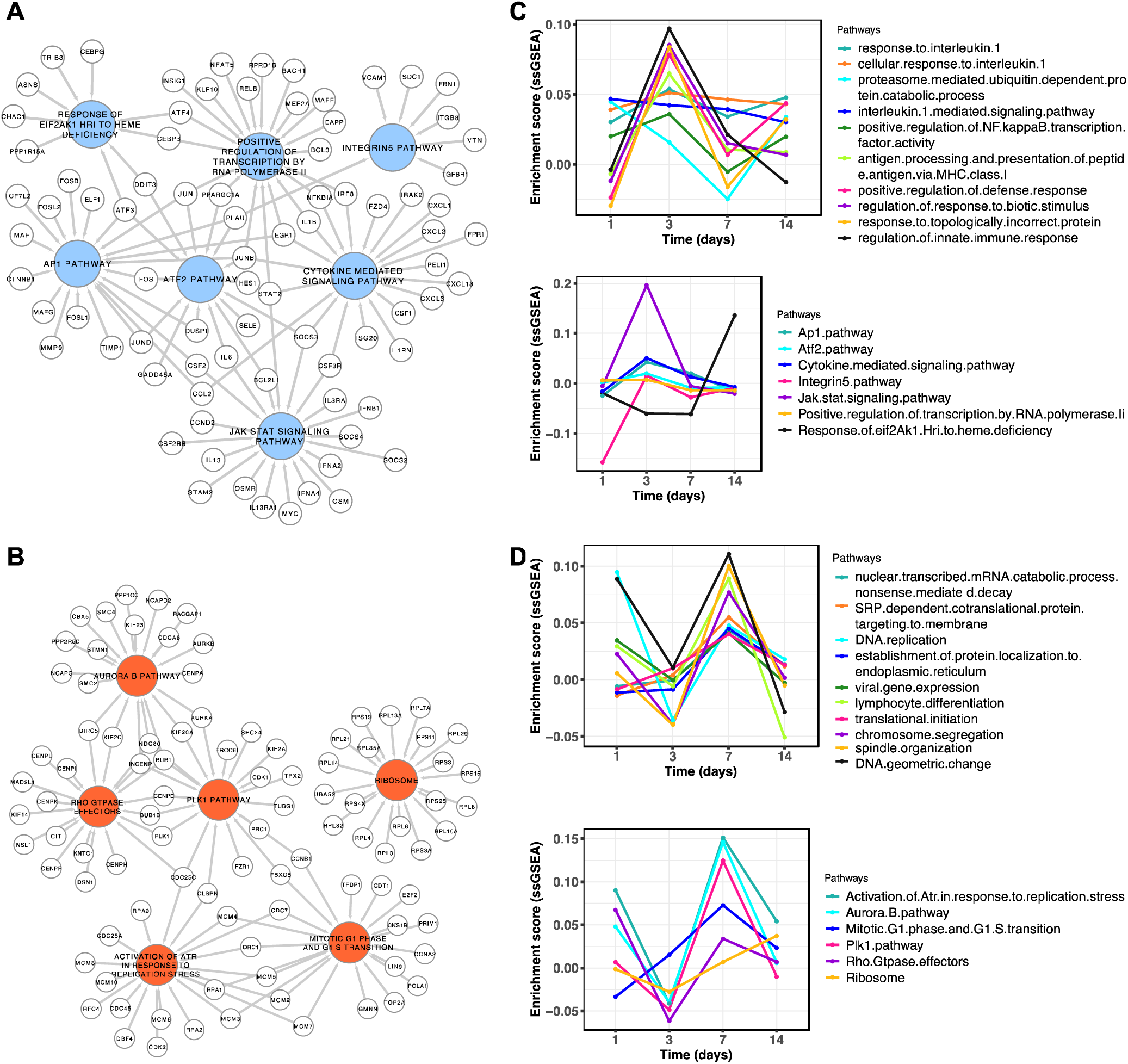
Dysregulated pathways or gene ontology terms induced by coronavirus infection and time-dependent host response dynamics in SARS-CoV-2. **A, B**, (**A**) Down- or (**B**) Up-regulated pathways enriched in the valid signatures and their example genes. Colored circles indicate pathways and small circles indicate genes involved in the connected pathways. For a clear visualization, no more than 20 genes in each pathway were shown. The networks were generated in Cytoscape (v 3.8.0). **C, D**, Dynamics of virus-induced biological processes in ferrets across three time points (in **C**, Top: down-regulated biological processes, and Bottom: down-regulated canonical pathways; in **D**, Top: up-regulated biological processes, and Bottom: up-regulated canonical pathways in the valid signatures) (study id: GSE147507). For each point, the enrichment score of pathway genes against infected samples was subtracted by the score of those genes against mock samples. ssGSEA with normalization across samples was applied.

More than one hundred suppressed genes were related to cellular responses to cytokine stimulus, inflammatory response, cytokine production, etc. (Figure 3A). This corroborates with other studies (Channappanavar *et al*. 2016; de Wit *et al*. 2016), which showed that SARS-CoV and MERS-CoV proteins interplay with the down-regulated JAK-STAT pathway in order to evade host immune response (Figure 3A). In addition to cytokine-related genes, other down-regulated genes include those involved in inflammatory mediators AP1 and ATF2 pathways (e.g., *JUN, FOS* and *CCL2*) (Figure 3A), and those related to IL1, IL4, IL6, IL10, and IL13 signaling. These interleukins were deficient in the early stage of SARS-CoV infection, caused by delayed type 1 interferon signaling, but then rapidly led to over-activated inflammatory cytokine storms (Channappanavar *et al*. 2016). These down-regulated genes were also identified by two recent studies of SARS-CoV-2 infection. Gordon *et al*. (Gordon *et al*. 2020) identified 332 high confidence SARS-CoV-2-human protein-protein interactions, among which 310 host genes were also found in our valid signatures. For instance, TBKBP1 (related to type 1 interferons production), TRIM59, and several other cytokine signaling related proteins bind to SARS-CoV-2 proteins (e.g. Nsp13 and Orf3a). Blanco-Melo *et al*. (Blanco-Melo *et al*. 2020) profiled the SARS-CoV-2 transcriptional signatures *in vitro* and *in vivo*, and observed a muted immune response that lacked robust induction of cytokines including type I and type III interferons compared with influenza A virus and human orthopneumovirus. These findings confirmed one common characteristic of MERS-CoV, SARS-CoV, and SARS-CoV-2 captured by our signatures: the suppression of cytokine production and signaling.

In addition to immune evasion, enrichment analysis of the down-regulated genes of our valid signatures suggested that viral infection inhibited host gene transcription (GO: Positive Regulation Of Transcription By RNA Polymerase II), probably *via* the viral Nsp-1 protein-induced host mRNA degradation (Narayanan *et al*. 2008). Host response to amino acid deficiency or unfolded protein pathways (REACTOME: Response of EIF2AK1 (HRI) to heme deficiency) were also suppressed (Figure 3A) by the virus in order to fully utilize the host cell machinery for its own replication. Moreover, the virus down-regulated the integrin pathway (Figure 3A), suggesting that it may disrupt intercellular surface interactions to prevent the infected cells from connecting with their neighbors. The importance of SARS-CoV-1/2 recognition by integrins was highlighted by Luan *et al*.(Luan *et al*. n.d.)

In terms of over-expressed genes, 33 ribosome genes were commonly up-regulated in our infection signatures (Figure 3B), suggesting the virus had already hijacked the host cell machinery for its own replication. The biological processes including mitotic cell cycle, nuclear division, and DNA replication were significantly over-represented. This might be related to Rho GTPase, PLK1, and Aurora B signaling activation, as implied by the up-regulated genes in our signatures (Figure 3B). The SARS-CoV-2 human protein interactome (Gordon *et al*. 2020, 2) revealed that Nsp7 binds to the host Rho A GTPase, agreeing with our infection signature where multiple Rho GTPase effectors were up-regulated, perhaps providing survival benefits to the virus (Van den Broeke, Jacob, and Favoreel 2014). The interactome also revealed a high overlap between the SARS-CoV-2 Nsp1-human protein interactome and proteins specific to the G1/S transition of mitotic cell cycle. Interestingly, coronavirus infection also promoted ATR signaling (Figure 3B), probably in response to DNA replication stress. This suggests a potential strategy to intervene with DNA replication stress induced by viral proteins, e.g., *via* the replication protein A complex (Xu *et al*. 2011).

After the first submission of this work, gene expression profiles induced by SARS-CoV-2 infection were released (GSE147507) (Blanco-Melo *et al*. 2020). We created several SARS-CoV-2 signatures from the *in vivo* SARS-CoV-2 infection model and found four signatures that could identify the drugs known to be effective against SARS-CoV-2 *in vitro* (Table S5). The dysregulated biological processes of SARS-CoV-2 signatures showed remarkable overlap with our valid 13 infection signatures derived from MERS-CoV and SARS-CoV, including suppressed cytokine signaling and ATF2 pathways, together with up-regulated mitotic cell cycle, PLK1, ATR and Rho GTPase pathways (Extended Data 5). To illustrate the dynamics of host response to SARS-CoV-2 infection *in vivo*, critical biological processes and pathways were analyzed across four time points within 14 days post-infection. In the early stages of infection (within 3 days post-infection), immune response processes (innate immune response, IL1 signaling, NF-kB, JAK-STAT) were stimulated (Figure 3C). These immune signaling pathways were suppressed from day 3 to day 7, then normalized to baseline at day 14, when the virus had been cleared from the body. In addition, three processes including proteasome-ubiquitin dependent protein catabolic process, host gene transcription by RNA polymerase II, and the EIF2AK1 response of heme deficiency remained inactive until day 7. In contrast, cell cycle related and viral replication processes were first suppressed, and then dramatically activated from day 3 to day 7 (Figure 3D). We also observed activation of non-sense mediated decay, ER localization, and G1-S transition from day 1 to day 7 (Figure 3D). The dynamics of virus-induced host gene expression changes suggested a tipping point of infection (day 3 in this model), where host cells were hijacked by the virus, resulting in immune suppression, host gene translation silencing, and activation of protein synthesis. These findings were also supported by further analysis, where infection signatures derived from the comparison of day 7 and day 3, rather than the comparison of day 3 and day 1, resulted in significant enrichment in positive drugs (p = 0.001, sRGES and EC_50_ correlation = 0.53, Table S5). This suggests that reversal of the hijacked gene expression changes between a specific time interval may be useful in designing novel antiviral therapeutics and strategies.

## Discussion

In this study, we investigated host gene expression changes induced by coronavirus (mainly SARS-CoV and MERS-CoV) infection to characterize the infection signatures for use in drug discovery for SARS-CoV-2. Out of 215 infection signatures, only 13 signatures could be used to accurately predict drug candidates that have been reported to be active against SARS-CoV or MERS-CoV. A large number of signatures did not provide useful information, and this could be due to variations in experimental models or conditions (such as virus incubation times). For example, samples using virus strains (dORF6, BatSRBD) in GSE4796 might be different from the wildtype SARS. Similarly, samples using ferret as the model for immune response investigation in study GSE22581 might require additional deconvolution of profiles to dissect the response in host cells. In study GSE79218, dysregulated pathways induced by MERS-CoV in human microvascular endothelial cells showed marked variations depending on the time post-infection (Figure S3E and F). Those disease signatures were not able to capture the biology of viral infection, thus failed to recover known drugs. These observations underscore the challenge of choosing appropriate experimental models and conditions (especially time points post-infection) in our bioinformatics approach to find repurposed drugs for SARS-CoV-2 (Kupferschmidt and Cohen 2020).

After the first submission of this work, gene expression profiles induced by SARS-CoV-2 infection were released. We then computed the SARS-CoV-2 signatures in A549, Calu3, and NHBE cells. NHBE and A549 cells with low viral load (MOI 0.2) did not show substantial transcriptional changes (less than 50 differentially expressed genes mapped to LINCS), partially due to their poor permissibility for SARS-CoV-2 (Hoffmann *et al*. 2020) and low viral load. A greater number of signature genes were identified in Calu3 and A549 cells with high viral load (MOI 2.0); however, the positive control drugs showed a virus-mimicking effect rather than a reversal of disease gene expression (Table S5). In Calu3 and A549 cells, only the 24 h post-infection profiles were available; however, other studies have shown that many infection signature genes have reverted back to normal at this time point (Figure S3). Considering the infection dynamics and varying cell permissibility, these signatures at 24 h might only capture the characteristics of host cell immune response against the virus, and not that of other processes such as viral replication. Indeed, the *in vivo* experiment with 14 days drug treatment performed in the same study (Hoffmann *et al*. 2020) led to signatures that could recover known drugs.

We developed a consensus score for each drug based on their potency to reverse the 13 valid infection signatures. The prediction was externally validated by a recent drug screen against SARS-CoV-2 (Jeon *et al*. 2020), and justified our approach based on reversal of host cell gene expression induced by SARS-CoV or MERS-CoV infection. Based on various criteria including predicted drug ranking and known MoA, we selected 10 drugs to test their antiviral efficacy and cytotoxicity in the Vero E6 cell line. We observed that four of these 10 drugs prevented CPE within 1 µM, but three of them were toxic at this concentration. We initially excluded only highly toxic compounds such as topoisomerase inhibitors, but did not attempt to evaluate their cytotoxicity since these candidates are FDA approved drugs or undergoing clinical trials, and therefore have established safety profiles.

The most potent candidate, methotrexate, is a chemotherapeutic agent at high doses and an immune suppressor at lower doses (used to treat rheumatoid arthritis (RA)). Methotrexate showed considerable *in vitro* antiviral effect and relatively low toxicity. The direct binding target of methotrexate, dihydrofolate reductase (DHFR), is an important enzyme in the folate metabolism pathway. DHRF is up-regulated in most of our valid signatures, together with the folate transporter SLC46A1 (Extended Data 3). Folate is required for the metabolism of several amino acids and is vital for nucleic acid synthesis, which could be utilized by the virus. From the LINCS drug profiles we found that methotrexate could down-regulate genes related to G2/M cell cycle (e.g. *MELK, KIF14* and *BIRC5*), and up-regulate genes related to cytokine signaling (e.g. *CCL2, CXCL2* and *NFKBIA*) (Figure 2D). This is congruent with previous observations that some interferon related pathways governing inflammation overlap with cancer (Musella *et al*. 2017). Recently, another RA drug, tocilizumab, an IL-6 antibody, was used to treat SARS-CoV-2 patients with severe symptoms to protect them from life-threatening lung damage caused by cytokine storms. Further analysis of drug-induced *in vivo* profiles (GEO accession: GSE56426 and GSE25160) revealed that methotrexate and tocilizumab could reverse several genes in our infection signatures that govern negative regulation of viral genome replication and viral life cycle (annotated with GO:0045071 and GO:1903901, Figure S6). Chloroquine has a lesser reversal effect on these genes, suggesting a different MoA from methotrexate and tocilizumab. Interestingly, most of these virus-blocking genes are also interferon-stimulated genes (ISGs) such as *ISG15* and *OAS1*. Thus, further testing of methotrexate in interferon sufficient cell lines compared with interferon deficient Vero cells could be of great interest.

The other three candidates are anti-cancer agents, namely bortezomib, nvp-bez235, and alvocidib. In cancer cells, proteasome inhibition by bortezomib induces ER stress and constitutive ER stress causes calcium release, resulting in apoptosis (Landowski *et al*. 2005). NVP-BEZ235 inhibits PI3K kinase and mTOR kinase in the PI3K/AKT/mTOR kinase signaling pathway, which may result in apoptosis of hyperactive cells and inhibition of growth in PI3K/mTOR-overexpressing cells (Netland *et al*. 2016). Alvocidib is a CDK inhibitor, acting through inhibition of *MCL1* expression that leads to apoptosis of cells (Arbour *et al*. 2008). In other independent drug repurposing efforts for anti-SARS-CoV-2, anti-cancer drugs like AXL receptor tyrosine kinase inhibitor Gilteritinib and CDK inhibitor Abemaciclib were also identified *in vitro* (Jeon *et al*. 2020; Weston *et al*. 2020). These anti-cancer drugs inhibit SARS-CoV-2 infection likely through targeting the hijacked host cells with hyperactive transcription and translation; however, it is a double-edged sword with good efficacy yet unfavorable cytotoxicity, especially for severely ill patients weakened by over-activation of the inflammatory system.

Importantly, the 13 valid infection signatures also provided a comprehensive understanding of the impact of coronavirus infection on the host response, especially the time-dependent dynamics of changes in biological processes, which may provide useful insights into designing clinically useful treatment strategies. These signatures captured several critical viral processes related not only to MERS-CoV and SARS-CoV infections, but also to SARS-CoV-2 infection, and thus might be applicable to studying future pathogens within this coronaviruses family. Rationally targeting these dysregulated pathways at specific time intervals may result in promising therapeutics against coronavirus infection.

Our study is restricted by the limitations of the LINCS datasets (e.g., limited coverage of the transcriptome). Another limitation is the use of drug-induced gene expression profiles derived from cancer cells that are different from the virus-infected cells used to generate the infection signatures. This may partially explain why antiviral drugs were not predicted as hits. Nevertheless, given the currently limited resources and the imperative need to find treatments for COVID-19, existing FDA-approved drugs that are identified, regardless of established MoA, should be further studied in infectious disease models.

In conclusion, open science initiatives have allowed us to leverage public resources to rationally predict drug candidates that might reverse coronavirus-induced transcriptomic changes. The unbiased search for drug candidates based on reversal of gene expression could offer an effective and rapid means to propose drug candidates for further experimental testing, even those that may have unexpected MoA. However, more layers of information such as toxicity, experimental validation conditions, and clinical applicability could be incorporated to find improved therapeutics. The predicted drug list and valid infection signatures resulting from our study may provide a starting point for researchers to further validate these and other candidates during this time of urgency.

## STAR Methods

### Computation of infection signatures

We obtained a total of 430 samples for “SARS-CoV” or “MERS-CoV” related data from ArrayExpress, Gene Expression Omnibus (GEO), and Sequence Read Archive (SRA). The metainformation of each sample was manually annotated, including virus strain, model, organism, and time point. The expression matrix for each microarray data was downloaded *via* the GEOquery R package. The matrix was further filtered by removing the probes with expression in only half of the samples. Expression values were normalized using quantile normalization, and log_2_ transformation was applied for each matrix. The probe values were collapsed based on Entrez Gene ID. The Significance Analysis of Microarrays (SAM) method was used to compute differentially expressed (DE) genes with criteria fold change > 1 and false discovery rate (FDR) < 0.05. Gene symbols of other organisms were converted to HUGO gene symbols. For RNA-Seq datasets, raw sequence data were downloaded from SRA and processed with the TOIL pipeline(Liu *et al*. 2019; Vivian *et al*. 2017). EdgeR was used to compute DE genes using the same criteria as used for microarray data. Gene ontology enrichment analysis of DE genes for each comparison was performed using the clusterprofiler R package. Further, gene set enrichment (ssGSEA) for each biological process was performed using ssGSEA method in the GSVA R package. For the infection group, we enumerated all the comparisons across all time points, and corresponding comparisons were performed in the mock group. The DE genes that were uniquely present in the infection group were selected for further analysis. We also compared DE genes between infection and mock groups at each time point, together with consistently dysregulated genes from the first to last time point.

### Computation of drug signatures

Drug gene expression profiles have been widely used in our previous studies. Briefly, a full matrix comprising 476,251 signatures and 22,268 genes including 978 landmark genes (as of September 2013) was downloaded from the LINCS website (https://clue.io). The metainformation of the signatures (for example, cell line, treatment duration, treatment concentration) was retrieved *via* LINCS Application Program Interfaces. The matrix and metadata are now available *via* GSE92742 in GEO. The signature derived from the comparison of gene expressions between the perturbagen- or vehicle control-treated samples represents gene expression changes upon treatment. We further downloaded the LINCS drug information from the Drug Repurposing Hub. Only small molecules with high quality gene expression profiles (is_gold=1, annotated in the metainformation), and which are listed in the Drug Repurposing Hub were further analyzed.

### Reversal correlation

The computation of Reversal of Gene Expression Score (RGES) and the summarization of RGES (to give the summarized RGES, or sRGES) were detailed elsewhere and recently implemented as a standalone R package (Zeng *et al*. 2019). In short, we quantified the reversal of disease gene expression as RGES, a measure modified from the connectivity score developed in other studies (Sirota et al. 2011; Subramanian et al. 2017). To compute RGES, we first rank genes based on their expression values in each drug profile. An enrichment score/s for each set of up- and down-regulated disease genes were computed separately using a Kolmogorov–Smirnov-like statistic, followed by merging of scores from both sets (up/down). The score is based on the extent to which the genes (up or down-regulated disease genes) are located at either the top or bottom of the ranked drug profile. One compound may have multiple available expression profiles because they have been tested in various cell lines, drug concentrations, treatment durations, or occationally different replicates, resulting in multiple RGES for one disease prediction. We set a reference condition (i.e., concentration of 10 µM, treatment duration of 24 h) and used a model to estimate a new RGES if the drug profile under the reference condition was not available. We summarized these scores as sRGES without weighting the cell lines. We considered predictions to be insignificant if the maximum of the absolute sRGES is < 0.25.

### Validation and selection of infection signatures

Drugs with known *in vitro* activity against two coronaviruses (i.e., SARS-CoV and MERS-CoV) served as positive controls to select valid infection signatures. Qualifying signatures should meet the following criteria: (1) derived from SARS-CoV or MERS-CoV infection experiments; (2) the number of differentially expressed genes was > 50 (mapped to LINCS); (3) the maximum absolute sRGES prediction was > 0.25; (4) the sRGES of positive drugs was enriched at the top (one side Wilcoxon rank-sum test p < 0.05, FDR < 0.25); (5) the sRGES and the average EC_50_ value of positive drugs were highly correlated (Spearman r >= 0.4, p < 0.05).

### Clustering of virus predictions

We downloaded the compiled virus-perturbed signatures from EnrichR (323 in total). Since the EnrichR dataset did not include any MERS-CoV signatures, we manually added the signatures of two MERS-CoV datasets (GSE79218, GSE79172, 8 in total) computed from the comparisons of separate infected groups and mock groups. Only virus signatures containing more than 50 LINCS landmark genes were selected. Each virus signature was queried against the LINCS library using the established pipeline. Only those signatures where the maximum of the absolute sRGES was > 0.25 were chosen for subsequent analysis. Viruses were clustered based on the sRGES scores using pvclust (Suzuki and Shimodaira 2006) (distance method: Spearman correlation, nboot = 100).

### Ritonavir correlation analysis

Ritonavir was originally developed as an inhibitor of HIV protease and is now often used at a low dose with other protease inhibitors to boost its antiviral effect. We found one RNA-Seq dataset (SRA: SRX4939022) for HepaRG cells treated with multiple compounds including ritonavir under multiple concentrations (ranging from 9 nM to 300 µM). We processed 90 profiles in the plate (2D_RG_PLATE2) consisting of 10 concentrations (each concentration has nine profiles). The log_2_ TPM (Transcripts Per Million) of each profile was subtracted by the median of log_2_ TPM of all DMSO-treated samples in this plate, resulting in one drug-induced gene expression profile. The Spearman correlation between the drug-induced gene expression and the disease gene expression used for the LINCS prediction was computed. A negative correlation means a reversal relationship.

### Visualization of infection signature reversal

To allow easier visualization of the selected drug candidates’ ability to reverse coronavirus-induced gene expression changes, we combined the 13 valid infection signatures into one meta signature, and summarized drug profiles from different experiments into one profile. Dysregulated genes were included into the meta infection signature if 25% quantile of log_2_ fold changes was < -1 or 75% quantile was > 2. For each drug, all profiles (z-scores, level 5) in L1000 measured at 10 µM were extracted (including different cell lines and treatment times). The value of each gene in the summarized profile was defined as the median of the head or tail 25% (depending on which absolute value is larger); if this quantile absolute value was > 1, we defined it as the median of the z-scores of this gene across all the profiles extracted. The matrix was composed of the meta signature genes and signatures of selected drug candidates. We also included the profiles of two drugs predicted as negative hits, and ordered the rows by the fold change of infection signature genes. A heatmap was used to visualize the effect of selected drugs in reversing virus-induced genes.

When visualizing the transcriptome datasets (for Figure S6) from GEO, we processed the gene expression matrix in the same manner as the disease signature mentioned under “Computation of infection signature”. The log2 fold change values of all genes were then converted to ranking percentages. Finally, a clustermap was computed for genes of our interest, colored in red (up-regulation) or blue (down-regulation).

### Enrichment Analysis

The processed compound transcriptome profile was categorized into up-or down-regulated genes, with a threshold of log_2_ fold change > 1 or < -1, respectively. Each group of genes was then submitted to Enrichr (E. Y. Chen *et al*. 2013) (https://amp.pharm.mssm.edu/Enrichr/) to compute the Gene Ontology (GO) enrichment analysis. GO terms with p-value < 0.05 and adjusted p-value < 0.05 were considered significant. We also used MSigDB (Subramanian *et al*. 2005) (https://www.gsea-msigdb.org/gsea/msigdb/index.jsp) to calculate the enriched GO (“C5”) and canonical pathways (“CP” under “C2”) enriched by the common dysregulated genes shown in more than half of the valid infection signatures.

### Cell culture, virus infection, and drug evaluation

Vero E6 cells [CRL:1586, ATCC] were grown in Eagle’s minimal essential medium (EMEM) supplemented with penicillin (100 units/ml), streptomycin (100 µg/ml), and 10% fetal bovine serum (FBS). SARS-CoV-2 (US_WA-1 isolate), the 3^rd^ passage in Vero E6 cells from the original CDC (Atlanta) material and sequence confirmed, was used throughout the study. The titer of the viral stock was 7.5 × 10^7^ 50% tissue culture infectious doses (TCID_50_)/ml. All experiments involving infectious virus were conducted at the University of Texas Medical Branch in an approved biosafety level 3 laboratory.

A slightly modified Vero E6-based standard micro-neutralization assay was used to rapidly evaluate the efficacy of predicted drugs against SARS-CoV-2 infection. Briefly, confluent Vero E6 cells grown in 96-wells microtiter plates were pre-treated with serially 2-folds diluted individual drugs for 3 h in the first instance, and 2 h in the repeat experiment, before infection with 100 infectious SARS-CoV-2 particles in 100 μl EMEM supplemented with 2% FBS. Vero E6 cells treated with similarly diluted dimethyl sulfoxide (DMSO) with or without virus were included as positive and negative controls, respectively.

After cultivation at 37 °C for 4 days, individual wells were observed under the microscope to determine virus-induced CPE and the effects of tested drugs. The efficacy of individual drugs was calculated and expressed as the lowest concentration capable of completely preventing virus-induced CPE in 100% of the wells. Toxicity to the treated cells was assessed by observing floating cells and altered morphology of adhered Vero E6 cells in wells under the microscope. All compounds were ordered from Selleckchem (USA) or Cayman Chemical (USA). All compounds were dissolved in 100% DMSO as 10 mM stock solutions and diluted in culture media.

### Software tools and statistical methods

All analyses were conducted in R (v3.5.1) or Python (v3.7) programming language. The ggplot2, pheatmap, and seaborn packages were used for data visualization. Student’s t-test was performed for normally distributed data and Wilcoxon rank-sum test was used for other types of data to compute the p-value.

### Data and code availability

Authors declare that all data used in this study are available within the article and its supplementary information files. Other specific files can be provided by the corresponding author upon reasonable request. The code is available at GitHub (https://github.com/Bin-Chen-Lab/wars).

## Author Contributions

B.C. conceived and supervised the study. J.X., R.S. and B.C. performed computational analyses with input from S.P. and E.C.. A.D. performed biological experiments supervised by C.T.K.T.. S.R. provided clinical insights, and T.D. prepared reagents. J.X., R.S., B.C., A.D., C.T.K.T wrote the manuscript with the input from all coauthors. M-S.C. provided feedback, reviewed and edited the manuscript.

## Acknowledgements

This research is supported by R01GM134307, K01 ES028047, and the MSU Global Impact Initiative. The content is solely the responsibility of the authors and does not necessarily represent the official views of sponsors. The authors would like to thank all researchers who shared their data publicly, making this project possible.

## References

Arbour, Nicole et al. 2008. “Mcl-1 Is a Key Regulator of Apoptosis during CNS Development and after DNA Damage.” The Journal of Neuroscience 28(24): 6068.

Bellani, Giacomo et al. 2016. “Epidemiology, Patterns of Care, and Mortality for Patients With Acute Respiratory Distress Syndrome in Intensive Care Units in 50 Countries.” JAMA 315(8): 788–800.

Blanco-Melo, Daniel et al. 2020. “SARS-CoV-2 Launches a Unique Transcriptional Signature from in Vitro, Ex Vivo, and in Vivo Systems.” bioRxiv: 2020.03.24.004655.

Caly, Leon et al. 2020. “The FDA-Approved Drug Ivermectin Inhibits the Replication of SARS-CoV-2 in Vitro.” Antiviral Research: 104787.

“CDC Cases and Updates.” https://www.cdc.gov/coronavirus/2019-ncov/cases-in-us.html.

Channappanavar, Rudragouda et al. 2016. “Dysregulated Type I Interferon and Inflammatory Monocyte-Macrophage Responses Cause Lethal Pneumonia in SARS-CoV-Infected Mice.” Cell Host & Microbe 19(2): 181–93.

Chen, Bin, Wei Wei, et al. 2017. “Computational Discovery of Niclosamide Ethanolamine, a Repurposed Drug Candidate That Reduces Growth of Hepatocellular Carcinoma Cells In Vitro and in Mice by Inhibiting Cell Division Cycle 37 Signaling.” Gastroenterology 152(8): 2022–36.

Chen, Bin, Li Ma, et al. 2017. “Reversal of Cancer Gene Expression Correlates with Drug Efficacy and Reveals Therapeutic Targets.” Nature Communications 8(1): 16022.

Chen, Edward Y. et al. 2013. “Enrichr: Interactive and Collaborative HTML5 Gene List Enrichment Analysis Tool.” BMC Bioinformatics 14(1): 128.

Chu, C M et al. 2004. “Role of Lopinavir/Ritonavir in the Treatment of SARS: Initial Virological and Clinical Findings.” Thorax 59(3): 252.

“COVID-19 Report from WHO.” https://www.who.int/emergencies/diseases/novel-coronavirus-2019/situation-reports/ (April 3, 2020).

Gordon, David E. et al. 2020. “A SARS-CoV-2-Human Protein-Protein Interaction Map Reveals Drug Targets and Potential Drug-Repurposing.” bioRxiv: 2020.03.22.002386.

Hoffmann, Markus et al. 2020. “SARS-CoV-2 Cell Entry Depends on ACE2 and TMPRSS2 and Is Blocked by a Clinically Proven Protease Inhibitor.” Cell 181(2): 271-280.e8.

Huang, Chaolin et al. 2020. “Clinical Features of Patients Infected with 2019 Novel Coronavirus in Wuhan, China.” The Lancet 395(10223): 497–506.

Jeon, Sangeun et al. 2020. “Identification of Antiviral Drug Candidates against SARS-CoV-2 from FDA-Approved Drugs.” bioRxiv: 2020.03.20.999730.

Kupferschmidt, Kai, and Jon Cohen. 2020. “Race to Find COVID-19 Treatments Accelerates.” Science 367(6485): 1412.

Landowski, Terry H. et al. 2005. “Mitochondrial-Mediated Disregulation of Ca^2+^ Is a Critical Determinant of Velcade (PS-341/Bortezomib) Cytotoxicity in Myeloma Cell Lines.” Cancer Research 65(9): 3828.

Lee, Bernard Kok Bang et al. 2017. “DeSigN: Connecting Gene Expression with Therapeutics for Drug Repurposing and Development.” BMC Genomics 18(1): 934.

Lim, Jaegyun et al. 2020. “Case of the Index Patient Who Caused Tertiary Transmission of Coronavirus Disease 2019 in Korea: The Application of Lopinavir/Ritonavir for the Treatment of COVID-19 Pneumonia Monitored by Quantitative RT-PCR.” J Korean Med Sci 35(6). https://doi.org/10.3346/jkms.2020.35.e79.

Liu, Ke et al. 2019. “Evaluating Cell Lines as Models for Metastatic Breast Cancer through Integrative Analysis of Genomic Data.” Nature Communications 10(1): 2138.

Lu, Roujian et al. 2020. “Genomic Characterisation and Epidemiology of 2019 Novel Coronavirus: Implications for Virus Origins and Receptor Binding.” Lancet (London, England) 395(10224): 565–74.

Luan, Junwen, Yue Lu, Shan Gao, and Leiliang Zhang. “A Potential Inhibitory Role for Integrin in the Receptor Targeting of SARS-CoV-2.” Journal of Infection. https://doi.org/10.1016/j.jinf.2020.03.046 (April 25, 2020).

Musella, Martina et al. 2017. “Type-I-Interferons in Infection and Cancer: Unanticipated Dynamics with Therapeutic Implications.” OncoImmunology 6(5): e1314424.

Narayanan, Krishna et al. 2008. “Severe Acute Respiratory Syndrome Coronavirus Nsp1 Suppresses Host Gene Expression, Including That of Type I Interferon, in Infected Cells.” Journal of Virology 82(9): 4471.

Netland, I. A. et al. 2016. “Dactolisib (NVP-BEZ235) Toxicity in Murine Brain Tumour Models.” BMC Cancer 16(1): 657.

van Noort, Vera et al. 2014. “Novel Drug Candidates for the Treatment of Metastatic Colorectal Cancer through Global Inverse Gene-Expression Profiling.” Cancer Research 74(20): 5690.

Peeri, Noah C et al. 2020. “The SARS, MERS and Novel Coronavirus (COVID-19) Epidemics, the Newest and Biggest Global Health Threats: What Lessons Have We Learned?” International Journal of Epidemiology (dyaa033). https://doi.org/10.1093/ije/dyaa033 (February 23, 2020).

Pessetto, Ziyan Y. et al. 2016. “In Silico and in Vitro Drug Screening Identifies New Therapeutic Approaches for Ewing Sarcoma.” Oncotarget 8(3). https://www.oncotarget.com/article/13385/text/.

Qu, Xiaoyan A., and Deepak K. Rajpal. 2012. “Applications of Connectivity Map in Drug Discovery and Development.” Drug Discovery Today 17(23): 1289–98.

Shen, Liang et al. 2019. “High-Throughput Screening and Identification of Potent Broad-Spectrum Inhibitors of Coronaviruses” ed. Tom Gallagher. Journal of Virology 93(12): e00023–19.

Sirota, Marina et al. 2011. “Discovery and Preclinical Validation of Drug Indications Using Compendia of Public Gene Expression Data.” Science Translational Medicine 3(96): 96ra77.

Subramanian, Aravind et al. 2005. “Gene Set Enrichment Analysis: A Knowledge-Based Approach for Interpreting Genome-Wide Expression Profiles.” Proceedings of the National Academy of Sciences 102(43): 15545.

———. 2017. “A Next Generation Connectivity Map: L1000 Platform and the First 1,000,000 Profiles.” Cell 171(6): 1437-1452.e17.

Suzuki, Ryota, and Hidetoshi Shimodaira. 2006. “Pvclust: An R Package for Assessing the Uncertainty in Hierarchical Clustering.” Bioinformatics 22(12): 1540–42.

Van den Broeke, Céline, Thary Jacob, and Herman W Favoreel. 2014. “Rho’ing in and out of Cells.” Small GTPases 5(1): e28318.

Vivian, John et al. 2017. “Toil Enables Reproducible, Open Source, Big Biomedical Data Analyses.” Nature Biotechnology 35(4): 314–16.

Wang, Manli et al. 2020. “Remdesivir and Chloroquine Effectively Inhibit the Recently Emerged Novel Coronavirus (2019-NCoV) in Vitro.” Cell Research. https://doi.org/10.1038/s41422-020-0282-0.

Weston, Stuart et al. 2020. “FDA Approved Drugs with Broad Anti-Coronaviral Activity Inhibit SARS-CoV-2 <em>in Vitro</Em>.” bioRxiv: 2020.03.25.008482.

de Wit, Emmie, Neeltje van Doremalen, Darryl Falzarano, and Vincent J. Munster. 2016. “SARS and MERS: Recent Insights into Emerging Coronaviruses.” Nature Reviews Microbiology 14(8): 523–34.

Wu, Chang-Jer et al. 2004. “Inhibition of Severe Acute Respiratory Syndrome Coronavirus Replication by Niclosamide.” Antimicrobial Agents and Chemotherapy 48(7): 2693.

Wu, Hongyu, Jinjiang Huang, Yang Zhong, and Qingshan Huang. 2017. “DrugSig: A Resource for Computational Drug Repositioning Utilizing Gene Expression Signatures.” PLOS ONE 12(5): e0177743.

Xu, Ling Hui, Mei Huang, Shou Guo Fang, and Ding Xiang Liu. 2011. “Coronavirus Infection Induces DNA Replication Stress Partly through Interaction of Its Nonstructural Protein 13 with the P125 Subunit of DNA Polymerase δ.” Journal of Biological Chemistry 286(45): 39546–59.

Zeng, Billy et al. 2019. “OCTAD: An Open Workplace for Virtually Screening Therapeutics Targeting Precise Cancer Patient Groups Using Gene Expression Features.” bioRxiv: 821546.

Zumla, Alimuddin et al. 2016. “Coronaviruses — Drug Discovery and Therapeutic Options.” Nature Reviews Drug Discovery 15(5): 327–47.

