## supplemental text for "Analysis of Infected Host Gene Expression Reveals Repurposed Drug Candidates and Time-Dependent Host Response Dynamics for COVID-19"

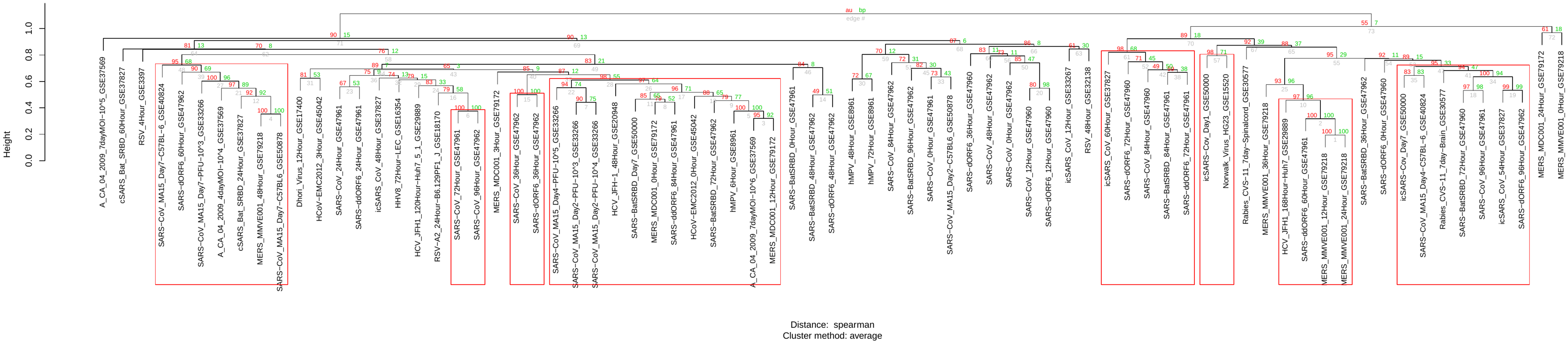
**Figure S1. Hierarchical clustering of virus infection signatures based on their drug prediction results.** Significant SARS/MERS clusters with approximately unbiased (AU) p-values no less than 95% were highlighted with red rectangles. Red and green numbers indicate AU and bootstrap probability (BP) p-values, respectively. Edge numbers are shown in grey. Among the 291 virus-induced signatures, only those that mapped to at least 50 LINCS landmark genes were used in this analysis.

**Table S1. Datasets used in this study for disease signature creation.**

| **S.**  **No.** | **Accession** | **Platform** | **Disease/**  **Infection** | **Organism** | **Model** | **Time points** | **Number of samples** | **PMID** |
| --- | --- | --- | --- | --- | --- | --- | --- | --- |
| 1 | GSE17400 | GPL570 | MOCK, DOHV and SARS | Homo sapiens | Calu-3 | 3 | 27 | 20090954 |
| 2 | GSE30589 | GPL570 | SARS and MOCK | Homo sapiens | Vero E6, Vero E6 DeltaE, MA-104 and MA-104 DeltaE | 4 | 33 | 22028656 |
| 3 | GSE45042 | GPL6480 | MOCK and EMC | Homo sapiens | Calu-3 | 6 | 33 | 23631916  24846384 |
| 4 | GSE47960 | GPL6480 | MOCK, SARS, H1N1, SARS-BatSRBD and SARS-dORF6 | Homo sapiens | HAE | 11 | 163 | 23935999 |
| 5 | GSE79218 | GPL13497 | MOCK and MERS | Homo sapiens | MMVE001 | 5 | 49 |  |
| 6 | GSE79172 | GPL13497 | MOCK and MERS | Homo sapiens | MDC001 | 5 | 29 | 28830941 |
| 7 | GSE22581 | GPL3738 | MOCK and SARS | Canis lupus | Ferret Lung | 3 | 9 | 21035159 |
| 8 | GSE36016 | GPL7202 | MOCK and SARS | Mus musculus | lung WT, lung IFNAR1 and lung STAT | 3 | 36 |  |
| 9 | GSE68820 | GPL7202 | MOCK and SARS | Mus musculus | lung wt and lung TLR3 | 3 | 52 | 26015500 |
| 10 | GSE30351 | GPL6480 | JFH1 | Homo sapiens | Huh7 | 2 | 7 |  |
| 11 | GSE71063 | GPL570 | HIV | Homo sapiens | patients | 2 | 40 | 26935044 |
| 12 | SRP166108 | Illumina HiSeq |  | Homo sapiens | HepaRG |  | 50 |  |

**Figure S2. Gene ontology enrichment of differentially expressed genes obtained from different time points in MOCK group of samples.** Gene ontology terms enriched in MOCK infected samples in **(A)** lung tissues without any knockout and **(B)** lung tissues with TLR3 knockout in GSE68820, and **(C)** MMVE001 cell lines in GSE79172. Terms enriched in MOCK infected samples are mostly cell cycle related. The size of the circles represents the ratio of genes in each process and color of the circle represents whether the genes are up- (red) or down- (green) regulated.

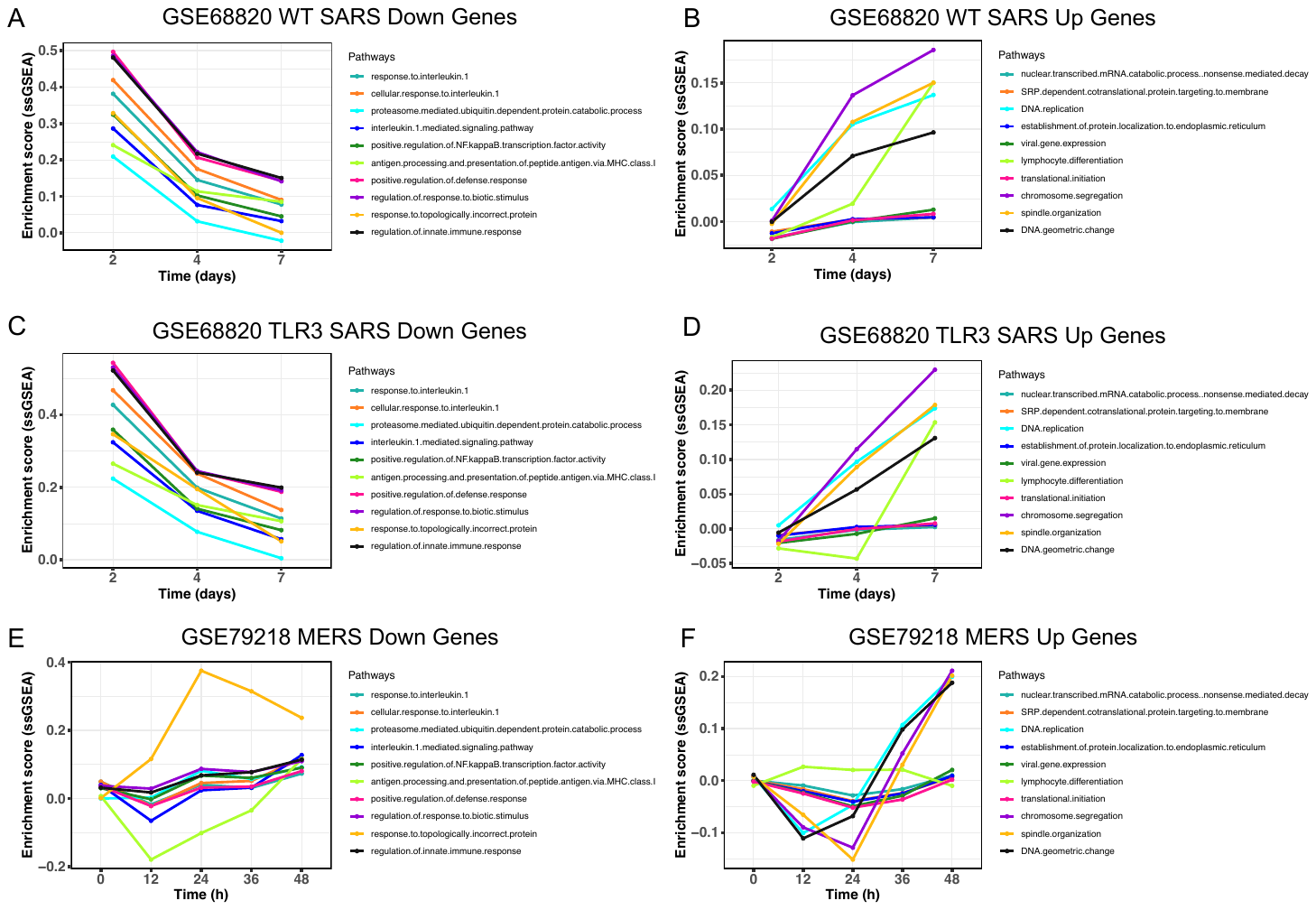

**Figure S3. Gene set enrichment of the biological biological processes at different time points.** Variations of gene set enrichment (ssGSEA) of biological processes obtained at different time points in down- **(A)** and up- **(B)** regulated genes induced by SARS-CoV infection in the wild type lung tissue. Similarly, variation of ssGSEA of biological processes obtained at different time points in down- **(C)** and up- **(D)** regulated genes induced by SARS-CoV infection in lung tissue with TLR3 mutant in GSE68820, and down- **(E)** and up- **(F)** regulated genes induced by MERS-CoV infection in MDC001 cell lines in GSE79172.

**Table S2. Positive control drugs with known activity against MERS-CoV/SARS-CoV/SARS-CoV-2.**

| **Name** | **MERS EC50 µM** | **SARS EC50 µM** | **COVID-19 EC50 µM** | **MoA** | **PMID** |
| --- | --- | --- | --- | --- | --- |
| amodiaquine | 6.21 | 1.27 | NA* | Antiparasitic agent | 32006468 |
| astemizole | 4.88 | 5.59 | NA | Neurotransmitter inhibitor | 32006468 |
| bisindolylmaleimide-ix | inhibition 74% @ 10uM | NA | NA | PKC inhibitor | 25653449 |
| bufalin | inhibition > 90% @ 10 nM | NA | NA | inhibit MERS-CoV entry by blocking clathrin-mediated endocytosis | 25653449 |
| chloroquine | 3 | NA | 1.13 | Endosomal acidification inhibitor | 32006468 |
| chlorpromazine | 4.9 | NA | NA | Neurotransmitter inhibitor | 32006468 |
| clomipramine | 9.33 | 13.23 | NA | Neurotransmitter inhibitor | 32006468 |
| dasatinib | 5.46 | 2.1 | NA | ABL1 inhibitor | 32006468 |
| disulfiram | NA | NA | NA | MERS-CoV PL-pro inhibitor | 32006468 |
| emetine | 0.34 | NA | NA | Inhibits RNA, DNA and protein synthesis | 32006468 |
| fluphenazine | 5.86 | 21.43 | NA | Neurotransmitter inhibitor | 32006468 |
| fluspirilene | 7.47 | 5.96 | NA | Neurotransmitter inhibitor | 32006468 |
| gemcitabine | 1.21 | 4.95 | NA | DNA metabolism inhibitor | 32006468 |
| GW-5074 | inhibition 52% @ 10uM | NA | NA | Raf inhibitor | 25653449 |
| imatinib | 17.68 | 9.82 | NA | ABL1 inhibitor | 32006468 |
| loperamide | 4.8 | 5.9 | NA | antidiarrheal opioid receptor agonist | 32006468 |
| lopinavir | 8 | 24.4 | NA | HIV-1 inhibitor | 32006468 |
| mefloquine | 7.41 | 15.55 | NA | Antiparasitic agent | 32006468 |
| mg-132 | NA | NA | NA | cys-protease m-calpain inhibition | 22787216 |
| monensin | 3.27 | NA | NA | Antibacterial | 32006468 |
| mycophenolate-mofetil | 1.54 | NA | NA | Immune suppressant, antineoplastic, antiviral | 32006468 |
| niclosamide | NA | 2 | NA | Antiparasitic agent | 15215127 |
| nitazoxanide | NA | NA | 2.12 | antiprotozoal, type I IFN inducer | 32020029 |
| ouabain | inhibition 70% @ 50 nM | NA | NA | inhibit MERS-CoV entry by blocking clathrin-mediated endocytosis | 25653449 |
| penciclovir | NA | NA | 95.96 | Inhibition of virus DNA synthesis | 32020029 |
| phenazopyridine | 1.93 | NA | NA | Analgesic | 32006468 |
| promethazine | 11.8 | 7.54 | NA | Neurotransmitter inhibitor | 32006468 |
| pyrvinium-pamoate | 1.84 | NA | NA | Anthelmintic | 32006468 |
| ribavirin | NA | NA | 109.5 | ribonucleic analog | 32020029 |
| ritonavir | 24.9 | NA | NA | HIV protease inhibitor | 31924756 |
| saracatinib | 2.9 | 2.4 | NA | Src family of tyrosine kinases inhibitor | 32006468 |
| SB-203580 | inhibition 45% @ 10uM | NA | NA | p38 MAPK inhibitor | 25653449 |
| selumetinib | inhibition >= 95% @ 10uM | NA | NA | MEK1, ERK1/2 inhibitor | 25653449 |
| sirolimus | inhibition 61% @ 10uM | NA | NA | MTOR inhibitor | 25653449 |
| tamoxifen | 10.11 | 92.88 | NA | Estrogen receptor inhibitor | 32006468 |
| terconazole | 12.2 | 15.32 | NA | Sterol metabolism inhibitor | 32006468 |
| thiothixene | 9.29 | 5.31 | NA | Neurotransmitter inhibitor | 32006468 |
| toremifene | 12.91 | 11.96 | NA | Estrogen receptor inhibitor | 32006468 |
| trametinib | inhibition >= 95% @ 10uM | NA | NA | MEK1/2 inhibitor | 25653449 |
| triflupromazine | 5.75 | 6.39 | NA | Neurotransmitter inhibitor | 32006468 |
| U-0126 | inhibition 51% @ 10uM | NA | NA | MEK1/2 inhibitor | 25653449 |

* “NA” indicates not available.

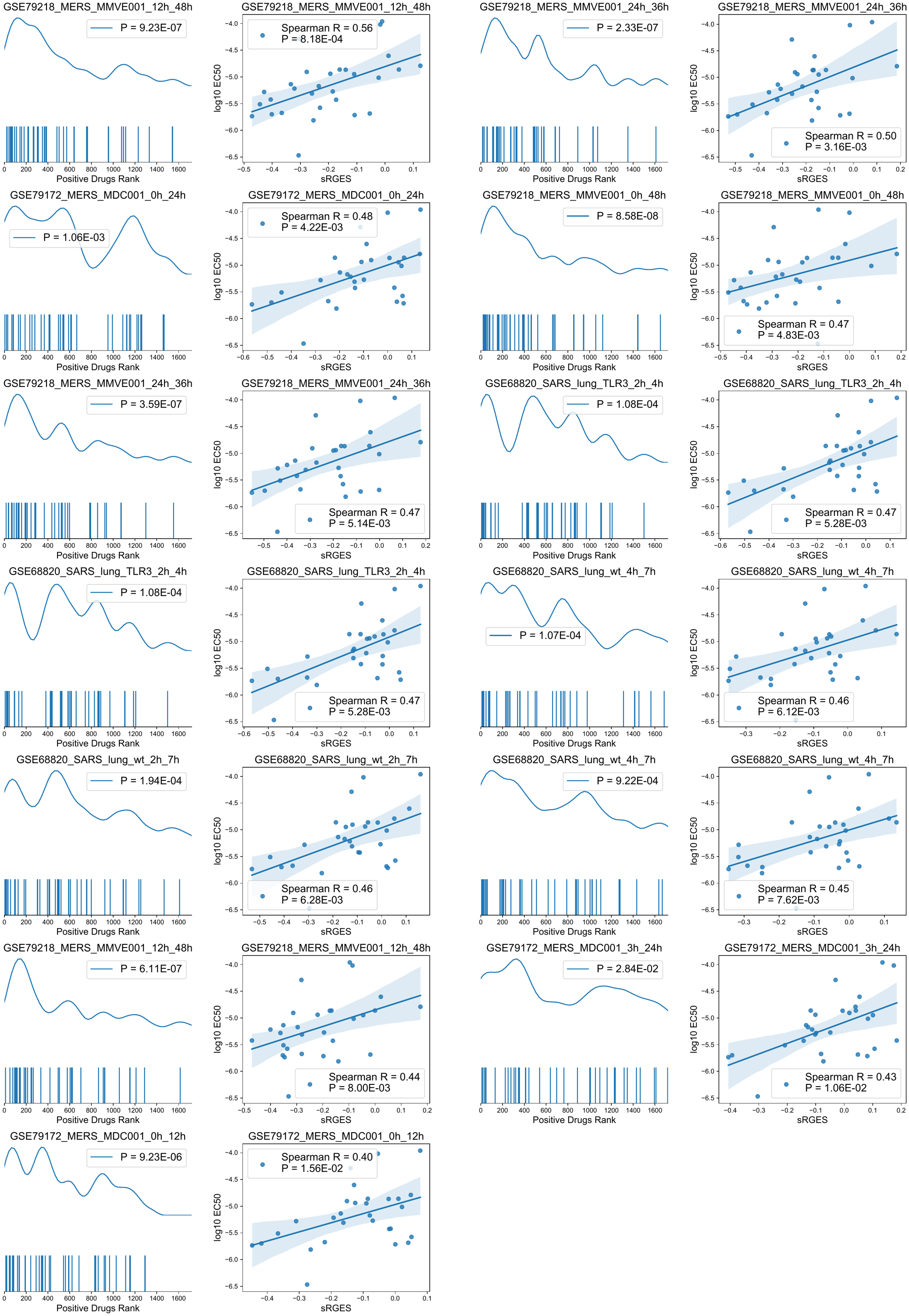

**Figure S4. SARS/MERS signatures validation using known active drugs (positive controls).** The first and third columns show enrichment density and barcode, with p values shown at upper right. The curve shows enrichment density, and each bar under the curve represents the rank of a positive drug among all the drugs in one prediction. The second and last columns list the correlation between sRGES and EC50 (in M, log10 transformed) of the positive controls, with Spearman R and p value shown at lower right. Each point indicates a positive control drug.

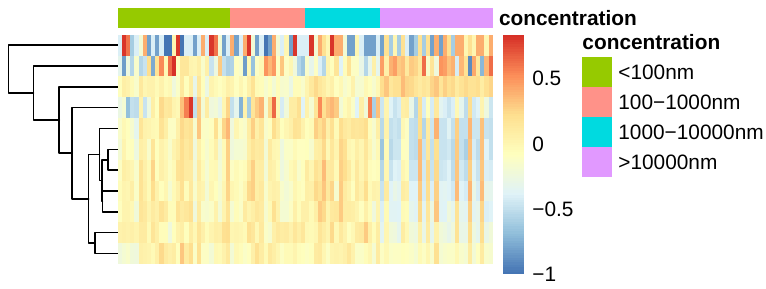

**Figure S5. Ritonavir profiles measured under different concentrations.** Each row indicates an infection signature, each column indicates a ritonavir profile under a different concentration annotated by the color bar on the right side. The heatmap shows the Spearman correlation coefficients between drug signatures and disease signatures. Blue means a stronger reversal effect.

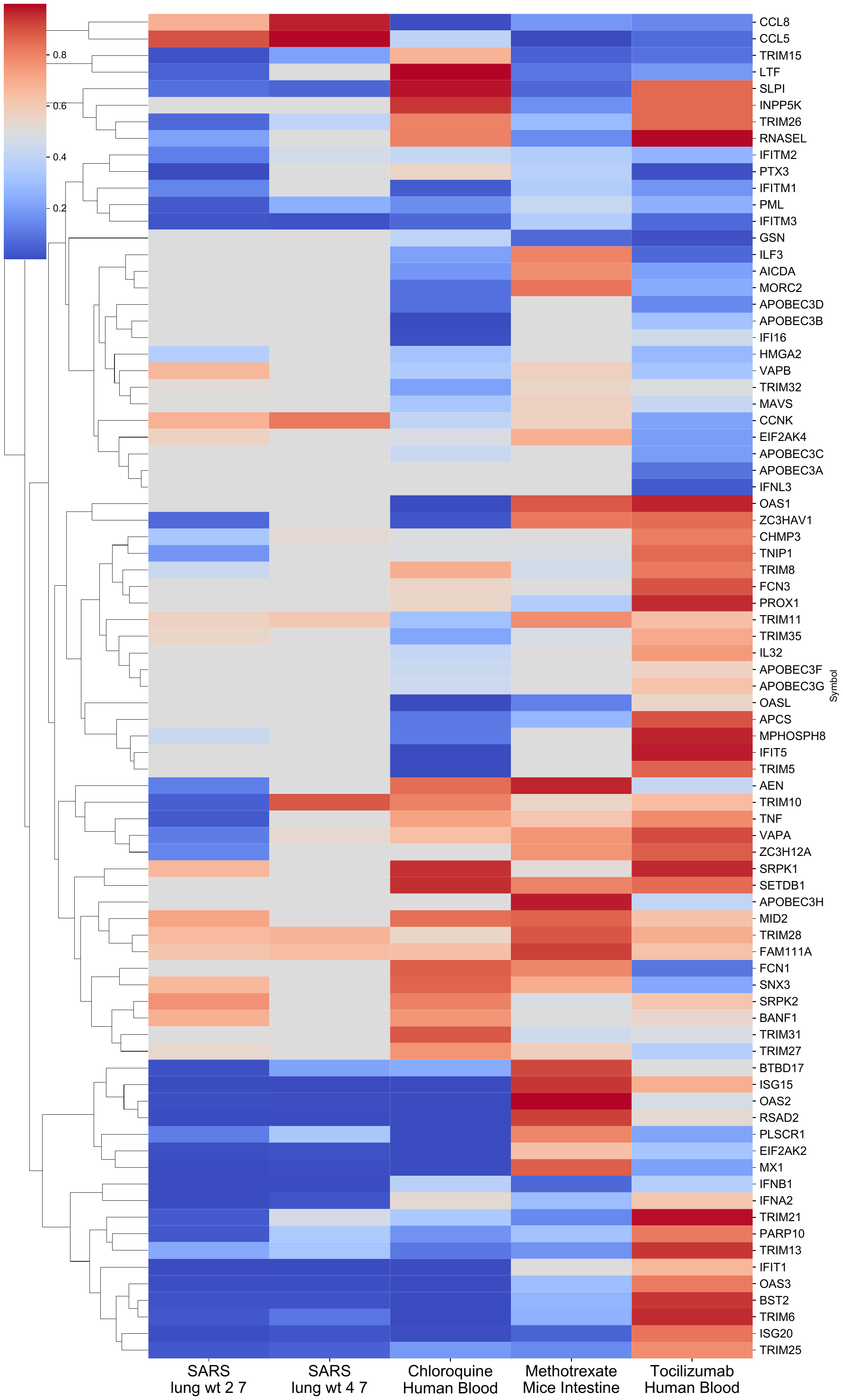

**Figure S6.** Heatmap showing the interferon-stimulated genes expression (rank percentage in the whole transcriptome) in infection (GSE68820), chloroquine (GSE71063), methotrexate (GSE56426) and tocilizumab (GSE25160) treated mice or patients. Red color indicates up-regulated genes and blue indicates down-regulated genes.

**Table S3. Antiviral prediction and other information of the ten selected candidates.**

| **Name** | **Median Rank (CoV)** | **Median Rank (MOCK)** | **Clinical Phase** | **Route of Administration** | **Targets** | **MoA** | **Indication** |
| --- | --- | --- | --- | --- | --- | --- | --- |
| Bortezomib | 3 | 23 | Launched | Intravenous | PSMA1-8  PSMB1-11  PSMD1-2  RELA | NFkB pathway inhibitor  Proteasome inhibitor | Multiple myeloma, mantle cell lymphoma |
| Puromycin | 18 | 128 | Launched | Oral | 60S ribosomal proteins | Protein synthesis inhibitor |  |
| Methotrexate | 37 | 356 | Launched | Oral | DHFR | Dihydrofolate reductase inhibitor | Gestational choriocarcinoma, hydatidiform mole, acute lymphoblastic leukemia, psoriasis, rheumatoid arthritis |
| Methylene blue | 57 | 153 | Launched | Intravenous | ACHE | Guanylyl cyclase inhibitor  nitric oxide production inhibitor | Methemoglobinemia |
| Tyloxapol | 106 | 293 | Launched | Inhalant | LPL  NFKB2 | NFkB pathway inhibitor |  |
| Nisoldipine | 108 | 245 | Launched | Oral | Voltage-dependent L-type calcium channels | Calcium channel blocker | Hypertension |
| Nvp-bez235 | 178 | 173 | Phase 2 | Oral | ATR  MTOR  PI3Ks | mTOR inhibitor  PI3K inhibitor |  |
| Fluvastatin | 283 | 362 | Launched | Oral | HMGCR | HMGCR inhibitor | Hypercholesterolemia, congenital heart defects |
| Alvocidib | 290 | 823 | Phase 2 | Intravenous | CDK1-2  CDK4-9  EGFR  PYGM | CDK inhibitor |  |
| Dasatinib | 378 | 1103 | Launched | Oral | ABL1/2  Src family kinases  EPHA2  KIT  PDGFRB  SRMS  STAT5B | Bcr-Abl kinase inhibitor  Ephrin inhibitor  KIT inhibitor  PDGFR tyrosine kinase receptor inhibitor  Src family inhibitor | Chronic myeloid leukemia, acute lymphoblastic leukemia |

**Table S4. Enrichment Analysis for Gene Ontology and Canonical Pathways of the Common Dysregulated Genes in the Valid Infection Signatures.**

| **Gene Set Name** | **# Genes in Gene Set (K)** | **# Genes in Overlap (k)** | **k/K** | **p-value** | **FDR q-value** | **Direction** |
| --- | --- | --- | --- | --- | --- | --- |
| GO RESPONSE TO CYTOKINE | 1210 | 162 | 0.134 | 1.23E-58 | 1.26E-54 | Down |
| GO REGULATION OF INTRACELLULAR SIGNAL TRANSDUCTION | 1863 | 202 | 0.108 | 4.57E-58 | 2.33E-54 | Down |
| GO LOCOMOTION | 1984 | 206 | 0.104 | 3.81E-56 | 1.30E-52 | Down |
| GO PROTEIN PHOSPHORYLATION | 1994 | 206 | 0.103 | 8.74E-56 | 2.23E-52 | Down |
| GO REGULATION OF CELL DEATH | 1748 | 187 | 0.107 | 1.13E-52 | 2.30E-49 | Down |
| GO CELL MOTILITY | 1758 | 187 | 0.106 | 2.67E-52 | 4.53E-49 | Down |
| GO RESPONSE TO OXYGEN CONTAINING COMPOUND | 1698 | 183 | 0.108 | 5.62E-52 | 8.19E-49 | Down |
| GO NEGATIVE REGULATION OF RESPONSE TO STIMULUS | 1703 | 183 | 0.108 | 8.69E-52 | 1.11E-48 | Down |
| GO REGULATION OF PROTEIN MODIFICATION PROCESS | 1880 | 190 | 0.101 | 8.32E-50 | 9.43E-47 | Down |
| GO DEFENSE RESPONSE | 1765 | 182 | 0.103 | 7.01E-49 | 7.15E-46 | Down |
| GO POSITIVE REGULATION OF SIGNALING | 1869 | 185 | 0.099 | 3.97E-47 | 3.67E-44 | Down |
| GO REGULATION OF PHOSPHORYLATION | 1625 | 171 | 0.105 | 5.03E-47 | 4.27E-44 | Down |
| GO REGULATION OF RESPONSE TO STRESS | 1559 | 167 | 0.107 | 6.31E-47 | 4.95E-44 | Down |
| GO REGULATION OF PHOSPHORUS METABOLIC PROCESS | 1823 | 182 | 0.100 | 7.29E-47 | 5.31E-44 | Down |
| GO POSITIVE REGULATION OF PROTEIN METABOLIC PROCESS | 1724 | 176 | 0.102 | 1.33E-46 | 9.01E-44 | Down |
| GO RESPONSE TO BIOTIC STIMULUS | 1567 | 166 | 0.106 | 5.40E-46 | 3.44E-43 | Down |
| GO POSITIVE REGULATION OF MULTICELLULAR ORGANISMAL PROCESS | 1825 | 180 | 0.099 | 1.38E-45 | 8.26E-43 | Down |
| GO REGULATION OF RESPONSE TO EXTERNAL STIMULUS | 1124 | 136 | 0.121 | 5.80E-44 | 3.29E-41 | Down |
| GO RESPONSE TO NITROGEN COMPOUND | 1135 | 136 | 0.120 | 1.76E-43 | 9.42E-41 | Down |
| GO REGULATION OF TRANSPORT | 1894 | 180 | 0.095 | 2.41E-43 | 1.23E-40 | Down |
| GO CELL ACTIVATION | 1442 | 154 | 0.107 | 5.03E-43 | 2.44E-40 | Down |
| GO REGULATION OF IMMUNE SYSTEM PROCESS | 1670 | 166 | 0.099 | 2.28E-42 | 1.06E-39 | Down |
| GO POSITIVE REGULATION OF MOLECULAR FUNCTION | 1781 | 171 | 0.096 | 1.09E-41 | 4.84E-39 | Down |
| GO POSITIVE REGULATION OF PROTEIN MODIFICATION PROCESS | 1247 | 140 | 0.112 | 1.65E-41 | 7.01E-39 | Down |
| GO WHOLE MEMBRANE | 1671 | 164 | 0.098 | 3.86E-41 | 1.57E-38 | Down |
| GO INFLAMMATORY RESPONSE | 765 | 108 | 0.141 | 4.11E-41 | 1.61E-38 | Down |
| GO POSITIVE REGULATION OF CELLULAR BIOSYNTHETIC PROCESS | 1988 | 181 | 0.091 | 4.91E-41 | 1.85E-38 | Down |
| GO MOLECULAR FUNCTION REGULATOR | 1812 | 171 | 0.094 | 1.04E-40 | 3.80E-38 | Down |
| GO SECRETION | 1680 | 163 | 0.097 | 2.96E-40 | 1.04E-37 | Down |
| GO BIOLOGICAL ADHESION | 1425 | 148 | 0.104 | 6.81E-40 | 2.31E-37 | Down |
| GO POSITIVE REGULATION OF DEVELOPMENTAL PROCESS | 1428 | 148 | 0.104 | 8.71E-40 | 2.86E-37 | Down |
| GO REGULATION OF CELL DIFFERENTIATION | 1881 | 173 | 0.092 | 9.86E-40 | 3.14E-37 | Down |
| GO POSITIVE REGULATION OF PHOSPHORUS METABOLIC PROCESS | 1146 | 131 | 0.114 | 1.17E-39 | 3.62E-37 | Down |
| GO REGULATION OF CELL POPULATION PROLIFERATION | 1716 | 163 | 0.095 | 4.19E-39 | 1.26E-36 | Down |
| GO RESPONSE TO ENDOGENOUS STIMULUS | 1721 | 163 | 0.095 | 6.01E-39 | 1.75E-36 | Down |
| GO CYTOKINE MEDIATED SIGNALING PATHWAY | 791 | 106 | 0.134 | 2.87E-38 | 8.14E-36 | Down |
| GO NEGATIVE REGULATION OF SIGNALING | 1427 | 145 | 0.102 | 5.45E-38 | 1.50E-35 | Down |
| GO POSITIVE REGULATION OF RNA METABOLIC PROCESS | 1710 | 160 | 0.094 | 1.43E-37 | 3.82E-35 | Down |
| GO CELLULAR RESPONSE TO OXYGEN CONTAINING COMPOUND | 1208 | 131 | 0.108 | 3.15E-37 | 8.24E-35 | Down |
| GO TUBE DEVELOPMENT | 1122 | 125 | 0.111 | 1.06E-36 | 2.71E-34 | Down |
| GO NEGATIVE REGULATION OF MULTICELLULAR ORGANISMAL PROCESS | 1318 | 136 | 0.103 | 2.53E-36 | 6.28E-34 | Down |
| REACTOME CYTOKINE SIGNALING IN IMMUNE SYSTEM | 858 | 109 | 0.127 | 3.53E-37 | 7.89E-34 | Down |
| GO POSITIVE REGULATION OF TRANSCRIPTION BY RNA POLYMERASE II | 1198 | 127 | 0.106 | 4.33E-35 | 1.03E-32 | Down |
| GO POSITIVE REGULATION OF NUCLEOBASE CONTAINING COMPOUND METABOLIC PROCESS | 1870 | 164 | 0.088 | 4.29E-35 | 1.03E-32 | Down |
| GO POSITIVE REGULATION OF CATALYTIC ACTIVITY | 1429 | 139 | 0.097 | 2.32E-34 | 5.36E-32 | Down |
| GO POSITIVE REGULATION OF INTRACELLULAR SIGNAL TRANSDUCTION | 1026 | 115 | 0.112 | 5.07E-34 | 1.15E-31 | Down |
| GO CYTOKINE PRODUCTION | 808 | 100 | 0.124 | 3.45E-33 | 7.63E-31 | Down |
| GO CELLULAR RESPONSE TO ENDOGENOUS STIMULUS | 1454 | 138 | 0.095 | 5.44E-33 | 1.18E-30 | Down |
| GO SIGNAL TRANSDUCTION BY PROTEIN PHOSPHORYLATION | 974 | 110 | 0.113 | 7.52E-33 | 1.60E-30 | Down |
| GO IMMUNE SYSTEM DEVELOPMENT | 990 | 110 | 0.111 | 3.21E-32 | 6.56E-30 | Down |
| GO REGULATION OF CELLULAR COMPONENT MOVEMENT | 1107 | 117 | 0.106 | 3.22E-32 | 6.56E-30 | Down |
| REACTOME SIGNALING BY INTERLEUKINS | 462 | 72 | 0.156 | 2.09E-30 | 2.33E-27 | Down |
| REACTOME INNATE IMMUNE SYSTEM | 1113 | 104 | 0.093 | 2.04E-24 | 1.52E-21 | Down |
| NABA MATRISOME | 1026 | 95 | 0.093 | 4.32E-22 | 2.41E-19 | Down |
| NABA MATRISOME ASSOCIATED | 751 | 79 | 0.105 | 7.17E-22 | 3.20E-19 | Down |
| KEGG CYTOKINE CYTOKINE RECEPTOR INTERACTION | 265 | 45 | 0.170 | 5.33E-21 | 1.98E-18 | Down |
| REACTOME SIGNALING BY RECEPTOR TYROSINE KINASES | 504 | 61 | 0.121 | 3.83E-20 | 1.22E-17 | Down |
| PID AP1 PATHWAY | 70 | 24 | 0.343 | 2.46E-19 | 6.86E-17 | Down |
| REACTOME INTERLEUKIN 4 AND INTERLEUKIN 13 SIGNALING | 111 | 28 | 0.252 | 2.68E-18 | 6.66E-16 | Down |
| REACTOME HEMOSTASIS | 678 | 63 | 0.093 | 4.27E-15 | 9.53E-13 | Down |
| REACTOME INTERLEUKIN 10 SIGNALING | 46 | 17 | 0.370 | 9.90E-15 | 1.90E-12 | Down |
| REACTOME NUCLEAR EVENTS KINASE AND TRANSCRIPTION FACTOR ACTIVATION | 61 | 19 | 0.312 | 1.02E-14 | 1.90E-12 | Down |
| KEGG MAPK SIGNALING PATHWAY | 267 | 37 | 0.139 | 1.37E-14 | 2.35E-12 | Down |
| REACTOME NGF STIMULATED TRANSCRIPTION | 39 | 15 | 0.385 | 1.85E-13 | 2.96E-11 | Down |
| KEGG TOLL LIKE RECEPTOR SIGNALING PATHWAY | 102 | 22 | 0.216 | 3.60E-13 | 5.36E-11 | Down |
| REACTOME EXTRACELLULAR MATRIX ORGANIZATION | 301 | 37 | 0.123 | 5.87E-13 | 8.19E-11 | Down |
| REACTOME TOLL LIKE RECEPTOR 9 TLR9 CASCADE | 95 | 21 | 0.221 | 7.40E-13 | 9.71E-11 | Down |
| REACTOME TOLL LIKE RECEPTOR CASCADES | 153 | 25 | 0.163 | 6.56E-12 | 8.13E-10 | Down |
| REACTOME NEUTROPHIL DEGRANULATION | 479 | 46 | 0.096 | 7.15E-12 | 8.40E-10 | Down |
| REACTOME SIGNALING BY NTRKS | 134 | 23 | 0.172 | 1.60E-11 | 1.79E-09 | Down |
| REACTOME DISEASE | 1470 | 93 | 0.063 | 1.97E-11 | 2.09E-09 | Down |
| REACTOME RESPONSE OF EIF2AK1 HRI TO HEME DEFICIENCY | 15 | 9 | 0.600 | 8.63E-11 | 8.37E-09 | Down |
| REACTOME TOLL LIKE RECEPTOR TLR1 TLR2 CASCADE | 97 | 19 | 0.196 | 8.61E-11 | 8.37E-09 | Down |
| NABA SECRETED FACTORS | 343 | 36 | 0.105 | 1.17E-10 | 1.09E-08 | Down |
| PID ATF2 PATHWAY | 59 | 15 | 0.254 | 1.67E-10 | 1.49E-08 | Down |
| REACTOME TOLL LIKE RECEPTOR 4 TLR4 CASCADE | 127 | 21 | 0.165 | 2.49E-10 | 2.14E-08 | Down |
| PID ERBB1 DOWNSTREAM PATHWAY | 105 | 19 | 0.181 | 3.60E-10 | 2.98E-08 | Down |
| REACTOME POST TRANSLATIONAL PROTEIN MODIFICATION | 1430 | 87 | 0.061 | 6.26E-10 | 4.99E-08 | Down |
| REACTOME MYD88 INDEPENDENT TLR4 CASCADE | 97 | 18 | 0.186 | 6.80E-10 | 5.06E-08 | Down |
| REACTOME TRANSPORT OF SMALL MOLECULES | 728 | 55 | 0.076 | 6.79E-10 | 5.06E-08 | Down |
| BIOCARTA IL1R PATHWAY | 31 | 11 | 0.355 | 8.86E-10 | 6.18E-08 | Down |
| PID IL6 7 PATHWAY | 47 | 13 | 0.277 | 8.84E-10 | 6.18E-08 | Down |
| REACTOME METABOLISM OF LIPIDS | 739 | 55 | 0.074 | 1.16E-09 | 7.87E-08 | Down |
| KEGG NOD LIKE RECEPTOR SIGNALING PATHWAY | 62 | 14 | 0.226 | 3.66E-09 | 2.41E-07 | Down |
| REACTOME TOLL LIKE RECEPTOR 10 TLR10 CASCADE | 84 | 16 | 0.191 | 4.03E-09 | 2.57E-07 | Down |
| KEGG PATHWAYS IN CANCER | 325 | 32 | 0.099 | 6.02E-09 | 3.73E-07 | Down |
| PID NFAT TFPATHWAY | 45 | 12 | 0.267 | 6.25E-09 | 3.77E-07 | Down |
| PID FRA PATHWAY | 37 | 11 | 0.297 | 7.58E-09 | 4.45E-07 | Down |
| REACTOME INTERLEUKIN 1 FAMILY SIGNALING | 139 | 20 | 0.144 | 8.02E-09 | 4.59E-07 | Down |
| REACTOME ION CHANNEL TRANSPORT | 183 | 23 | 0.126 | 8.93E-09 | 4.98E-07 | Down |
| REACTOME OVARIAN TUMOR DOMAIN PROTEASES | 38 | 11 | 0.290 | 1.04E-08 | 5.51E-07 | Down |
| KEGG JAK STAT SIGNALING PATHWAY | 155 | 21 | 0.136 | 1.02E-08 | 5.51E-07 | Down |
| PID INTEGRIN5 PATHWAY | 17 | 8 | 0.471 | 1.29E-08 | 6.70E-07 | Down |
| REACTOME CHEMOKINE RECEPTORS BIND CHEMOKINES | 58 | 13 | 0.224 | 1.46E-08 | 7.39E-07 | Down |
| REACTOME DEVELOPMENTAL BIOLOGY | 1131 | 70 | 0.062 | 1.59E-08 | 7.87E-07 | Down |
| REACTOME CLASS A 1 RHODOPSIN LIKE RECEPTORS | 331 | 31 | 0.094 | 3.31E-08 | 1.61E-06 | Down |
| REACTOME GPCR LIGAND BINDING | 463 | 38 | 0.082 | 3.50E-08 | 1.66E-06 | Down |
| REACTOME GLYCOSAMINOGLYCAN METABOLISM | 124 | 18 | 0.145 | 3.86E-08 | 1.79E-06 | Down |
| REACTOME PEPTIDE LIGAND BINDING RECEPTORS | 198 | 23 | 0.116 | 3.99E-08 | 1.82E-06 | Down |
| REACTOME INTERLEUKIN 1 SIGNALING | 102 | 16 | 0.157 | 7.19E-08 | 3.21E-06 | Down |
| REACTOME CELL CYCLE | 674 | 246 | 0.365 | 1.04E-167 | 2.32E-164 | Up |
| GO CELL CYCLE | 1864 | 374 | 0.201 | 1.54E-158 | 1.57E-154 | Up |
| REACTOME CELL CYCLE MITOTIC | 560 | 207 | 0.370 | 1.09E-141 | 1.21E-138 | Up |
| GO CELL CYCLE PROCESS | 1399 | 310 | 0.222 | 2.59E-142 | 1.32E-138 | Up |
| GO MITOTIC CELL CYCLE | 1030 | 258 | 0.251 | 7.89E-131 | 2.68E-127 | Up |
| GO CHROMOSOME ORGANIZATION | 1223 | 266 | 0.218 | 5.83E-119 | 1.49E-115 | Up |
| GO CHROMOSOME | 1730 | 292 | 0.169 | 2.08E-101 | 4.24E-98 | Up |
| GO CELL DIVISION | 598 | 172 | 0.288 | 9.59E-97 | 1.63E-93 | Up |
| GO DNA METABOLIC PROCESS | 925 | 200 | 0.216 | 5.63E-88 | 8.19E-85 | Up |
| REACTOME M PHASE | 416 | 138 | 0.332 | 1.30E-86 | 9.65E-84 | Up |
| REACTOME CELL CYCLE CHECKPOINTS | 291 | 117 | 0.402 | 1.54E-84 | 8.62E-82 | Up |
| GO CHROMOSOMAL REGION | 349 | 126 | 0.361 | 3.15E-84 | 4.02E-81 | Up |
| GO CHROMOSOME SEGREGATION | 324 | 122 | 0.377 | 4.26E-84 | 4.82E-81 | Up |
| GO CELL CYCLE PHASE TRANSITION | 637 | 163 | 0.256 | 4.24E-83 | 4.32E-80 | Up |
| GO ORGANELLE FISSION | 450 | 139 | 0.309 | 1.44E-82 | 1.33E-79 | Up |
| GO REGULATION OF CELL CYCLE | 1224 | 222 | 0.181 | 3.35E-82 | 2.84E-79 | Up |
| GO CELLULAR RESPONSE TO DNA DAMAGE STIMULUS | 856 | 182 | 0.213 | 1.29E-78 | 1.01E-75 | Up |
| GO DNA REPLICATION | 267 | 105 | 0.393 | 1.04E-74 | 7.60E-72 | Up |
| REACTOME MITOTIC PROMETAPHASE | 203 | 93 | 0.458 | 1.11E-73 | 4.97E-71 | Up |
| GO REGULATION OF CELL CYCLE PROCESS | 790 | 169 | 0.214 | 2.49E-73 | 1.69E-70 | Up |
| GO DNA REPAIR | 551 | 141 | 0.256 | 8.61E-72 | 5.48E-69 | Up |
| GO NUCLEAR CHROMOSOME SEGREGATION | 263 | 101 | 0.384 | 1.15E-70 | 6.87E-68 | Up |
| GO CONDENSED CHROMOSOME | 224 | 94 | 0.420 | 4.43E-70 | 2.51E-67 | Up |
| GO MICROTUBULE CYTOSKELETON | 1220 | 203 | 0.166 | 3.28E-68 | 1.76E-65 | Up |
| GO REGULATION OF MITOTIC CELL CYCLE | 664 | 146 | 0.220 | 6.52E-65 | 3.32E-62 | Up |
| GO DNA CONFORMATION CHANGE | 322 | 104 | 0.323 | 5.55E-64 | 2.69E-61 | Up |
| GO MITOCHONDRION | 1576 | 226 | 0.143 | 8.14E-64 | 3.77E-61 | Up |
| GO MITOTIC NUCLEAR DIVISION | 264 | 95 | 0.360 | 1.95E-63 | 8.64E-61 | Up |
| GO CHROMOSOME CENTROMERIC REGION | 193 | 82 | 0.425 | 9.18E-62 | 3.90E-59 | Up |
| GO DNA DEPENDENT DNA REPLICATION | 146 | 73 | 0.500 | 1.50E-61 | 6.12E-59 | Up |
| GO SISTER CHROMATID SEGREGATION | 189 | 81 | 0.429 | 2.17E-61 | 8.51E-59 | Up |
| GO SPINDLE | 355 | 103 | 0.290 | 2.94E-58 | 1.11E-55 | Up |
| REACTOME MITOTIC METAPHASE AND ANAPHASE | 235 | 85 | 0.362 | 4.68E-57 | 1.74E-54 | Up |
| REACTOME RESOLUTION OF SISTER CHROMATID COHESION | 126 | 65 | 0.516 | 4.46E-56 | 1.42E-53 | Up |
| GO RIBONUCLEOTIDE BINDING | 1893 | 236 | 0.125 | 2.79E-55 | 1.01E-52 | Up |
| GO REGULATION OF CELL CYCLE PHASE TRANSITION | 482 | 115 | 0.239 | 4.31E-55 | 1.52E-52 | Up |
| GO PROTEIN CONTAINING COMPLEX ASSEMBLY | 1782 | 226 | 0.127 | 3.80E-54 | 1.29E-51 | Up |
| GO CELLULAR PROTEIN CONTAINING COMPLEX ASSEMBLY | 1036 | 167 | 0.161 | 6.66E-54 | 2.19E-51 | Up |
| GO DRUG BINDING | 1724 | 221 | 0.128 | 1.05E-53 | 3.33E-51 | Up |
| GO MICROTUBULE BASED PROCESS | 752 | 141 | 0.188 | 1.32E-53 | 4.08E-51 | Up |
| GO MITOTIC SISTER CHROMATID SEGREGATION | 151 | 68 | 0.450 | 1.73E-53 | 5.19E-51 | Up |
| REACTOME RHO GTPASE EFFECTORS | 324 | 94 | 0.290 | 3.17E-53 | 8.84E-51 | Up |
| GO MICROTUBULE ORGANIZING CENTER | 759 | 141 | 0.186 | 4.29E-53 | 1.25E-50 | Up |
| GO ADENYL NUCLEOTIDE BINDING | 1541 | 205 | 0.133 | 2.72E-52 | 7.69E-50 | Up |
| GO MICROTUBULE CYTOSKELETON ORGANIZATION | 553 | 119 | 0.215 | 7.45E-52 | 2.05E-49 | Up |
| REACTOME SIGNALING BY RHO GTPASES | 454 | 108 | 0.238 | 1.24E-51 | 3.08E-49 | Up |
| REACTOME MITOTIC SPINDLE CHECKPOINT | 110 | 58 | 0.527 | 6.80E-51 | 1.38E-48 | Up |
| REACTOME RHO GTPASES ACTIVATE FORMINS | 140 | 64 | 0.457 | 6.32E-51 | 1.38E-48 | Up |
| GO SMALL MOLECULE METABOLIC PROCESS | 1962 | 233 | 0.119 | 8.65E-51 | 2.32E-48 | Up |
| REACTOME DNA REPAIR | 331 | 92 | 0.278 | 2.48E-50 | 4.61E-48 | Up |
| REACTOME SEPARATION OF SISTER CHROMATIDS | 190 | 72 | 0.379 | 3.84E-50 | 6.60E-48 | Up |
| GO MEIOTIC CELL CYCLE | 245 | 80 | 0.327 | 8.20E-50 | 2.14E-47 | Up |
| GO SUPRAMOLECULAR COMPLEX | 1285 | 180 | 0.140 | 5.20E-49 | 1.33E-46 | Up |
| GO CONDENSED CHROMOSOME CENTROMERIC REGION | 118 | 58 | 0.492 | 1.42E-48 | 3.53E-46 | Up |
| GO CENTROSOME | 579 | 117 | 0.202 | 5.37E-48 | 1.30E-45 | Up |
| GO DNA RECOMBINATION | 288 | 84 | 0.292 | 7.38E-48 | 1.75E-45 | Up |
| REACTOME CHROMOSOME MAINTENANCE | 117 | 57 | 0.487 | 1.79E-47 | 2.85E-45 | Up |
| GO ORGANONITROGEN COMPOUND BIOSYNTHETIC PROCESS | 1859 | 220 | 0.118 | 1.35E-47 | 3.12E-45 | Up |
| GO CATALYTIC COMPLEX | 1355 | 183 | 0.135 | 1.64E-47 | 3.72E-45 | Up |
| GO NUCLEAR CHROMOSOME | 1273 | 174 | 0.137 | 7.31E-46 | 1.62E-43 | Up |
| GO KINETOCHORE | 135 | 59 | 0.437 | 1.28E-45 | 2.77E-43 | Up |
| GO REGULATION OF ORGANELLE ORGANIZATION | 1267 | 172 | 0.136 | 6.38E-45 | 1.36E-42 | Up |
| GO MEIOTIC CELL CYCLE PROCESS | 187 | 67 | 0.358 | 6.98E-45 | 1.45E-42 | Up |
| REACTOME CELLULAR RESPONSES TO EXTERNAL STIMULI | 612 | 115 | 0.188 | 7.23E-44 | 1.08E-41 | Up |
| GO HYDROLASE ACTIVITY ACTING ON ACID ANHYDRIDES | 863 | 137 | 0.159 | 2.08E-43 | 4.24E-41 | Up |
| REACTOME DNA DOUBLE STRAND BREAK REPAIR | 166 | 62 | 0.374 | 6.86E-43 | 9.56E-41 | Up |
| KEGG CELL CYCLE | 124 | 52 | 0.419 | 3.12E-39 | 4.10E-37 | Up |
| REACTOME S PHASE | 161 | 58 | 0.360 | 3.59E-39 | 4.45E-37 | Up |
| REACTOME HOMOLOGY DIRECTED REPAIR | 138 | 54 | 0.391 | 8.90E-39 | 1.05E-36 | Up |
| REACTOME DNA REPLICATION | 127 | 52 | 0.409 | 1.37E-38 | 1.53E-36 | Up |
| REACTOME DISEASE | 1470 | 171 | 0.116 | 7.41E-36 | 7.87E-34 | Up |
| REACTOME G2 M CHECKPOINTS | 168 | 56 | 0.333 | 9.07E-36 | 9.20E-34 | Up |
| REACTOME MITOTIC G1 PHASE AND G1 S TRANSITION | 149 | 53 | 0.356 | 1.41E-35 | 1.37E-33 | Up |
| REACTOME DNA STRAND ELONGATION | 32 | 28 | 0.875 | 2.51E-35 | 2.33E-33 | Up |
| PID PLK1 PATHWAY | 46 | 31 | 0.674 | 1.46E-32 | 1.30E-30 | Up |
| REACTOME TRANSLATION | 295 | 67 | 0.227 | 4.21E-31 | 3.61E-29 | Up |
| REACTOME INFECTIOUS DISEASE | 804 | 113 | 0.141 | 5.81E-31 | 4.80E-29 | Up |
| REACTOME METABOLISM OF AMINO ACIDS AND DERIVATIVES | 374 | 75 | 0.201 | 7.32E-31 | 5.84E-29 | Up |
| KEGG DNA REPLICATION | 36 | 27 | 0.750 | 1.36E-30 | 1.04E-28 | Up |
| REACTOME POST TRANSLATIONAL PROTEIN MODIFICATION | 1430 | 158 | 0.111 | 1.44E-30 | 1.07E-28 | Up |
| REACTOME MITOTIC G2 G2 M PHASES | 200 | 55 | 0.275 | 2.72E-30 | 1.96E-28 | Up |
| REACTOME HDR THROUGH HOMOLOGOUS RECOMBINATION HRR | 67 | 34 | 0.508 | 1.25E-29 | 8.74E-28 | Up |
| REACTOME DEPOSITION OF NEW CENPA CONTAINING NUCLEOSOMES AT THE CENTROMERE | 73 | 35 | 0.480 | 2.37E-29 | 1.61E-27 | Up |
| REACTOME ACTIVATION OF THE PRE REPLICATIVE COMPLEX | 33 | 25 | 0.758 | 1.31E-28 | 8.59E-27 | Up |
| REACTOME METABOLISM OF RNA | 672 | 98 | 0.146 | 3.08E-28 | 1.97E-26 | Up |
| REACTOME TELOMERE MAINTENANCE | 90 | 36 | 0.400 | 1.05E-26 | 6.51E-25 | Up |
| REACTOME SRP DEPENDENT COTRANSLATIONAL PROTEIN TARGETING TO MEMBRANE | 113 | 39 | 0.345 | 5.67E-26 | 3.42E-24 | Up |
| REACTOME INFLUENZA INFECTION | 156 | 44 | 0.282 | 5.26E-25 | 3.09E-23 | Up |
| REACTOME ACTIVATION OF ATR IN RESPONSE TO REPLICATION STRESS | 37 | 24 | 0.649 | 6.92E-25 | 3.96E-23 | Up |
| REACTOME EUKARYOTIC TRANSLATION ELONGATION | 94 | 35 | 0.372 | 9.55E-25 | 5.33E-23 | Up |
| REACTOME ORGANELLE BIOGENESIS AND MAINTENANCE | 296 | 59 | 0.199 | 2.09E-24 | 1.14E-22 | Up |
| REACTOME MEIOSIS | 119 | 38 | 0.319 | 6.05E-24 | 3.22E-22 | Up |
| REACTOME EUKARYOTIC TRANSLATION INITIATION | 120 | 38 | 0.317 | 8.51E-24 | 4.42E-22 | Up |
| KEGG RIBOSOME | 88 | 33 | 0.375 | 1.58E-23 | 7.99E-22 | Up |
| REACTOME RESPONSE OF EIF2AK4 GCN2 TO AMINO ACID DEFICIENCY | 102 | 35 | 0.343 | 2.34E-23 | 1.16E-21 | Up |
| REACTOME RNA POLYMERASE II TRANSCRIPTION | 1375 | 139 | 0.101 | 3.20E-23 | 1.55E-21 | Up |
| REACTOME SELENOAMINO ACID METABOLISM | 118 | 37 | 0.314 | 4.88E-23 | 2.32E-21 | Up |
| REACTOME BASE EXCISION REPAIR | 91 | 33 | 0.363 | 5.51E-23 | 2.56E-21 | Up |
| PID AURORA B PATHWAY | 39 | 23 | 0.590 | 1.63E-22 | 7.44E-21 | Up |
| REACTOME TRANSCRIPTIONAL REGULATION BY TP53 | 365 | 63 | 0.173 | 1.87E-22 | 8.33E-21 | Up |

**Table S5. Positive control drugs prediction derived from SARS-CoV-2-induced transcriptional data.**

| **Infection Signature** | **Comparison** | **Host** | **P^*1^ (Enrichment)** | **FDR (Enrichment)** | **SpearmanR^*1^ (sRGES vs EC_50_)** | **P^*1^ (SpearmanR)** | **# of Genes Mapped** |
| --- | --- | --- | --- | --- | --- | --- | --- |
| Series11_Series13_ferret_SARS-CoV-2_d3-d14_all_gene^*2^ | Day14 vs Day3 | Ferret in vivo | 7.86E-05 | 0.001 | 0.499 | 0.003 | 619 |
| Series11_Series12_ferret_SARS-CoV-2_d3-d7_all_gene | Day7 vs Day3 | Ferret in vivo | 0.001 | 0.005 | 0.527 | 0.002 | 301 |
| Series10_Series13_ferret_SARS-CoV-2_d1-d14_all_gene | Day14 vs Day1 | Ferret in vivo | 0.001 | 0.005 | 0.383 | 0.020 | 655 |
| Series12_ferret_SARS-CoV-2_Ctl_d7_all_gene | Virus vs control; Day7 | Ferret in vivo | 0.006 | 0.016 | 0.471 | 0.005 | 334 |
| Series10_Series12_ferret_SARS-CoV-2_d1-d7_all_gene | Day7 vs Day1 | Ferret in vivo | 0.005 | 0.016 | 0.444 | 0.008 | 346 |
| Series6_A549_ACE2_SARS-CoV-2_DE_gene | Virus vs control | A549 cell line | 0.064 | 0.149 | -0.370 | 0.976 | 68 |
| Series13_ferret_SARS-CoV-2_Ctl_d14_all_gene | Virus vs control; Day14 | Ferret in vivo | 0.225 | 0.450 | 0.108 | 0.289 | 92 |
| Series12_Series13_ferret_SARS-CoV-2_d7-d14_all_gene | Day14 vs Day7 | Ferret in vivo | 0.273 | 0.478 | 0.370 | 0.024 | 585 |
| Series14_ferret_SARS-CoV-2_Ctl_d3_all_gene | Virus vs control; Day3 | Ferret in vivo | 1.000 | 1.000 | 0.002 | 0.496 | 263 |
| Series11_ferret_SARS-CoV-2_Ctl_d3_all_gene | Virus vs control; Day3 | Ferret in vivo | 1.000 | 1.000 | -0.159 | 0.794 | 288 |
| Series10_Series11_ferret_SARS-CoV-2_d1-d3_all_gene | Day3 vs Day1 | Ferret in vivo | 0.980 | 1.000 | -0.246 | 0.901 | 269 |
| Series10_ferret_SARS-CoV-2_d1_all_gene | Virus vs control; Day1 | Ferret in vivo | 0.986 | 1.000 | -0.299 | 0.942 | 387 |
| Series5_A549_SARS-CoV-2_DE_gene | Virus vs control | A549 cell line | 0.972 | 1.000 | -0.481 | 0.996 | 127 |
| Series7_Calu3_SARS-CoV-2_DE_gene | Virus vs control | Calu3 cell line | 0.984 | 1.000 | -0.515 | 0.998 | 138 |
| Series11_Series13_ferret_Ctl_d3-d14_all_gene | Day14 vs Day3; control only | Ferret in vivo | 0.025 |  | 0.553 | 0.001 | 593 |
| Series10_Series13_ferret_Ctl_d1-d14_all_gene | Day14 vs Day1; control only | Ferret in vivo | 0.000 |  | 0.390 | 0.018 | 585 |
| Series12_Series13_ferret_Ctl_d7-d14_all_gene | Day14 vs Day7; control only | Ferret in vivo | 0.001 |  | 0.382 | 0.020 | 636 |
| Series10_Series11_ferret_Ctl_d1-d3_all_gene | Day3 vs Day1; control only | Ferret in vivo | 0.086 |  | 0.156 | 0.210 | 362 |
| Series11_Series12_ferret_Ctl_d3-d7_all_gene | Day7 vs Day3; control only | Ferret in vivo | 0.999 |  | 0.099 | 0.305 | 273 |
| Series10_Series12_ferret_Ctl_d1-d7_all_gene | Day7 vs Day1; control only | Ferret in vivo | 0.964 |  | -0.157 | 0.792 | 386 |

^*1^ One-sided p values. The alternative hypothesis was that the positive controls enriched at top and sRGES positively correlated with efficacy.

^*2^ Valid signatures were highlighted in red.
